# Chromatin accessibility–transcription coupling is dynamically reconfigured during *Arabidopsis* embryo maturation

**DOI:** 10.64898/2026.06.23.733948

**Authors:** Daesong Jeong, Ilha Lee, Chulmin Park

## Abstract

Chromatin accessibility (CA) is widely profiled alongside transcription because it is generally associated with transcriptional activity. How directly the two layers are linked remains unclear. We addressed this in *Arabidopsis thaliana* embryos. We profiled CA and transcription across the torpedo, bent-cotyledon, and mature-green stages using stage-resolved nicking enzyme-assisted sequencing (NicE-seq) and nanopore RNA sequencing. To test whether their relationship is genetically regulated, we also analyzed *ld-1*, a mutant of *LUMINIDEPENDENS* (*LD*) linked to global transcriptional repression. Transcriptomes formed discrete stage-specific programmes, but CA did not. Accessibility at both promoters and transcription-factor motifs largely followed a common developmental trajectory that peaked at the bent-cotyledon stage and declined thereafter, rather than mirroring module-specific expression patterns. Loss of LD preserved promoter architecture but extensively remodelled motif accessibility, increasing its concordance with module-specific expression timing and strengthening CA–transcription coupling. Chromatin accessibility and transcription are subject to partly independent regulation during embryo maturation. The ability of *LD* loss to strengthen accessibility–transcription concordance demonstrates that their coupling is a regulated rather than fixed property of developmental chromatin.

## Introduction

Embryogenesis is a universal developmental process in which a single cell gives rise to a multicellular body plan through coordinated cell division, fate specification, and morphogenesis. In seed plants, embryogenesis also includes a maturation phase that prepares the embryo for dormancy and subsequent germination (Santos-Mendoza *et al*., 2008). This phase is accompanied by extensive chromatin reorganization. Cotyledon nuclei become smaller and exhibit increased chromatin condensation (van Zanten *et al*., 2011). DNA methylation patterns are dynamically reconfigured during seed maturation and germination (Bouyer *et al*., 2017; Kawakatsu *et al*., 2017). Histone variants also contribute to seed chromatin states. H3.3 is deposited at regulatory regions linked to post-embryonic developmental competence, whereas H2B.8 accumulates in mature embryos and is associated with chromatin compaction (Jiang *et al*., 2020; Buttress *et al*., 2022; Zhao *et al*., 2022). Despite these changes, how the chromatin accessibility (CA) landscape is reorganized during embryo maturation remains incompletely understood (Sullivan *et al*., 2019; Pei *et al*., 2023; Zhao *et al*., 2023; Zheng *et al*., 2025).

Accessible chromatin marks genomic regions permissive for transcription factor (TF) binding and cis-regulatory function. Combined CA and transcriptome profiling is therefore widely used to study developmental gene regulation. In animals, accessibility profiling has revealed dynamic remodeling of cis-regulatory elements during embryogenesis (Lu *et al*., 2016; Gao *et al*., 2018). In plants, similar approaches have been applied to roots, leaves, meristems, and flowers (Sijacic *et al*., 2018; Dorrity *et al*., 2021). In these studies, CA and transcription often show only partial concordance. Furthermore, many genes undergo substantial expression changes without corresponding CA alterations (Klemm *et al*., 2019; Kiani *et al*., 2022; Fillot & Mazza, 2025). Similarly, in *Arabidopsis* ecotypes, many differentially accessible regions are not associated with changes in nearby gene expression (Alexandre *et al*., 2018). For example, in shade-responsive seedlings, rapid transcriptional responses are often accompanied by little or no change in CA (Paulišić *et al*., 2025). Together, these observations suggest that the relationship between CA and transcription is context dependent. However, how CA–transcription coupling changes during developmental transitions, and whether this relationship can be genetically reconfigured, remain open questions in plants.

*Arabidopsis* provides a tractable system for addressing this question. Accessible chromatin regions in *Arabidopsis* are strongly enriched near promoters, with more than 75% located within 2 kb of the nearest transcription start site (Sijacic *et al*., 2018; Bubb & Deal, 2020). In addition, differences in CA between cell types are largely quantitative rather than qualitative (Sijacic *et al*., 2018). Together, these features suggest that regulatory information in *Arabidopsis* is concentrated predominantly in promoter-proximal regions. This compact regulatory architecture contrasts with large-genome species such as maize, where distal cis-regulatory elements and long-range enhancer–promoter interactions are widespread (Ricci *et al*., 2019). *Arabidopsis* therefore provides an opportunity to examine how tightly promoter accessibility tracks transcription during embryo maturation at genome-wide and motif-resolved scales.

Seed-related CA studies are beginning to emerge in plants. Regulatory DNA dynamics have been profiled in maturing *Arabidopsis* siliques (Sullivan *et al*., 2019), and CA together with transcriptome changes has been analyzed across embryo development in hexaploid wheat (Zhao *et al*., 2023). The wheat study reported broad concordance between accessibility and transcription, accompanied by maturation-associated chromatin changes including reduced accessibility and increased H3K27me3 (Zhao *et al*., 2023). CA has also been examined during wheat grain and maize seed development (Pei *et al*., 2023; Zheng *et al*., 2025). However, stage-resolved CA dynamics during *Arabidopsis* zygotic embryogenesis remain largely unexplored.

*Arabidopsis* embryogenesis proceeds through a series of morphologically defined stages, including globular, heart, torpedo, bent-cotyledon, and mature-green stages (Mansfield & Briarty, 1991; O’Neill *et al*., 2019). These stages define four successive developmental transitions: T1, globular to heart; T2, heart to torpedo; T3, torpedo to bent-cotyledon; and T4, bent-cotyledon to mature-green. Embryo development is governed by well-characterized transcriptional programs controlling early patterning and late maturation, including *WUSCHEL-RELATED HOMEOBOX* (*WOX*)-, *AUXIN RESPONSE FACTOR* (*ARF*)-, *CUP-SHAPED COTYLEDON* (*CUC*)-, and *CLASS III HOMEODOMAIN-LEUCINE ZIPPER* (*HD-ZIP III*)-dependent morphogenesis and the LAFL (*LEAFY COTYLEDON1* (*LEC1*), *ABSCISIC ACID INSENSITIVE3* (*ABI3*), *FUSCA3* (*FUS3*), and *LEC2*) maturation network (Baud *et al*., 2008; Holdsworth *et al*., 2008; Santos-Mendoza *et al*., 2008; Palovaara *et al*., 2017; Kao *et al*., 2021). In parallel, epigenetic states are progressively reorganized during embryo maturation, including increased CHH methylation at transposable elements (TEs) (Bouyer *et al*., 2017; Kawakatsu *et al*., 2017). However, the CA dynamics of individual TE families remain poorly characterized. Together, these features make *Arabidopsis* embryo maturation an informative system for examining how CA and transcription are coordinated during developmental transitions.

Most plant CA studies have relied on ATAC-seq or DNase-based approaches, which require freshly isolated nuclei from sufficient amounts of unfixed tissue (Bubb & Deal, 2020). These requirements are difficult to satisfy in *Arabidopsis* embryos, which must be individually dissected and pooled. Nicking enzyme assisted sequencing (NicE-seq) uses a nicking enzyme-based labeling strategy compatible with fixed, low-input material without requiring nuclear isolation or transposition (Ponnaluri *et al*., 2017; Chin *et al*., 2020; Vishnu *et al*., 2021). Despite these advantages, NicE-seq has not been applied to stage-resolved profiling of *Arabidopsis* embryos.

Here, we combined stage-resolved NicE-seq with matched RNA-seq to profile *Arabidopsis* embryos across mid-to-late embryogenesis. Core analyses focused on torpedo, bent-cotyledon, and mature-green stages to define genome-wide CA dynamics and their relationship to transcription during embryo maturation. To test whether limited CA–transcription concordance reflects a fixed chromatin property or a genetically regulated state, we additionally analyzed the *ld-1* loss-of-function mutant of *LUMINIDEPENDENS* (*LD*), which has recently been linked to transcriptional repression, premature transcription termination, and chromatin modification (He *et al*., 2003; Liu *et al*., 2007; Mateo-Bonmatí *et al*., 2024; Bergis-Ser *et al*., 2025).

## Materials and Methods

### Plant materials and growth conditions

*Arabidopsis thaliana* Columbia-0 (Col-0) was used as the wild-type (WT) control, and *ld-1* has been described previously (Rédei, 1962).

Seeds were sown on half-strength Murashige and Skoog (MS) medium (Duchefa, Haarlem, the Netherlands) supplemented with 1% sucrose and 1% plant agar (Duchefa). Seeds were stratified at 4 °C for 3 days to promote uniform germination. Seedlings were grown at 22 °C under long-day conditions (16 h light/8 h dark) using HO18-G3 lights containing white, blue, red, and far-red wavelengths, with a light intensity of 120–160 µmol m⁻² s⁻¹.

### Embryo staging

Developing embryos were staged morphologically under a stereomicroscope according to standard *Arabidopsis* criteria (Sullivan *et al*., 2019), using silique position on the main inflorescence as an additional guide. Under our conditions, heart-, torpedo-, bent-cotyledon-, and mature-green-stage embryos were collected primarily from siliques 9–11, 13–15, 17–19, and 21–23, respectively, counted from the youngest fertilized silique.

### Transcriptome profiling and analysis

Total RNA was extracted per biological replicate from 50 heart-, 30 torpedo-, 10 bent-cotyledon-, and 10 mature-green-stage embryos. Embryos were dissected in 20% RNAlater, washed three times, and transferred to 100 µL TRIzol (minimizing RNAlater carryover, <10 µL residual), incubated at 60 °C for 30 min, with bent-cotyledon and mature-green embryos additionally homogenized using a micro-pestle. RNA was purified with the Direct-zol RNA Microprep kit (Zymo Research). Full-length cDNA was generated from 4 ng total RNA using Smart-seq3 (Hagemann-Jensen *et al*., 2020). Libraries were prepared with the Native Barcoding Kit 24 V14 (Oxford Nanopore Technologies) and sequenced on a MinION Mk1B with an R10.4.1 flow cell; raw signal was basecalled with MinKNOW v25.09.16.

Raw Nanopore reads were processed with a UMI-aware workflow for Smart-seq3 libraries: UMIs were extracted as the 8-nt sequence adjacent to the template-switch oligo anchor, and reads were processed with pychopper v2.7.10 (edlib mode, PCS109 configuration, Smart-seq3 primer definitions) (Hagemann-Jensen *et al*., 2020), retaining full-length and rescued reads of ≥200 nt and mean quality ≥7. Spliced alignment to the *Arabidopsis thaliana* TAIR10 genome (Araport11 annotation) used minimap2 v2.28 (-ax splice -uf -k14 --secondary=no -L --cs=long -G 50k --junc-bonus 12), followed by samtools v1.19.2 (Li, 2018; Danecek *et al*., 2021). Reads were assigned to exons with HTSeq-count v2.0.5 (-s yes -t exon -i gene_id) (Anders *et al*., 2015) and deduplicated with UMI-tools (--per-gene --method=directional --edit-distance-threshold=2) (Smith *et al*., 2017).

Count matrices were analysed in R with DESeq2 v1.42.0 (additive genotype + stage design; size factors by poscounts; variance-stabilized transformation [VST] for PCA, clustering, and heatmaps) (Love *et al*., 2014). Throughout, lowly expressed genes were filtered with edgeR v4.0.16 filterByExpr (min.count = 5, min.total.count = 10) (Robinson *et al*., 2010). For module discovery, stage-mean VST values were gene-wise z-scored and k-means clustered (k = 6, 8, 10; set.seed(1), nstart = 50), with modules ordered by the stage of peak mean z-score. Differential expression between consecutive stages was tested per genotype with DESeq2 (Wald test, poscounts), calling genes at adjusted P < 0.05. GO Biological Process enrichment (clusterProfiler::enrichGO, org.At.tair.db) and GSEA (gseGO, genes ranked by the Wald statistic) used the expressed gene set as background, retaining terms at adjusted P < 0.05. Module × stage-transition DE overlap was tested by one-sided Fisher exact test against the expressed gene set (Benjamini–Hochberg per panel).

Promoter motif enrichment on module and DE gene sets used promoter regions (500 bp upstream to 200 bp downstream of the TSS) and monaLisa v1.8.1 calcBinnedMotifEnrR against JASPAR 2024 plant profiles (min.score = 7), testing foreground versus a GC-matched expressed-gene background (Machlab *et al*., 2022; Rauluseviciute *et al*., 2024). The position weight matrix (PWM) set was supplemented with custom consensus PWMs for embryo-relevant families absent from JASPAR (B3_RY, NF-Y, WOX, TCP_CL2, GRAS_SCR), and redundant PWMs (correlation > 0.90) were removed. TF families were normalised to a canonical form and resolved into motif-consensus-aware subfamily bins (Display_Bin; e.g., ERF into DREB, ERF_GCC, ERF_other; 1R-MYB into 1R-MYB_telo, 1R-MYB_GATA, 1R-MYB_other). Enrichment was called at adjusted P < 0.01, excluding gene sets with < 20 genes.

### Chromatin accessibility profiling and analysis (NicE-seq)

NicE-seq libraries were prepared using the enzyme-only, one-pot UniNicE-seq workflow as described (Ponnaluri *et al*., 2017; Chin *et al*., 2020; Vishnu *et al*., 2021), using 50 heart-, 30 torpedo-, 10 bent-cotyledon-, and 10 mature-green-stage embryos per replicate. Briefly, accessible chromatin was labelled by Nt.CviPII nicking and DNA polymerase I incorporation of biotinylated nucleotides; proteins were then digested in the same reaction, DNA was further fragmented by a second Nt.CviPII digestion, and labelled DNA was captured on streptavidin beads, end-repaired, adaptor-ligated, and PCR-amplified. Libraries were sequenced on an Illumina NovaSeq X.

Reads were trimmed with Trim Galore v0.6.11 (--quality 20 --length 20; --paired with explicit adapters where applicable) (Martin, 2011) and aligned to the *Arabidopsis thaliana* TAIR10 genome with Bowtie 2 v2.5.4 (end-to-end sensitive mode), followed by samtools v1.19.2 (Langmead & Salzberg, 2012; Danecek *et al*., 2021). Nuclear alignments with MAPQ ≥ 30 were retained (excluding unmapped, secondary, QC-fail, and supplementary reads), and duplicates were removed with samtools markdup. Normalized BigWig tracks were generated with deepTools v3.5.5 bamCoverage (--binSize 10 --minMappingQuality 30 --ignoreDuplicates; effective genome size 119,146,348) (Ramírez *et al*., 2016); reads-per-genomic-content (RPGC) normalization was used for metaprofiles, heatmaps, and BDI, CPM tracks for supplementary visualization, and SPMR tracks for MACS3 signal. Where depth-matched comparisons were required, BAMs were downsampled with sambamba v1.0.1 (Tarasov *et al*., 2015), except for count-based differential testing.

Broad accessibility peaks were called with MACS3 v3.0.2 (callpeak -f BAMPE -g 1.2e8 -q 0.05 -B --SPMR --nomodel --broad --broad-cutoff 0.1; single-end mode where applicable) (Zhang *et al*., 2008). Accessibility was quantified with Rsubread v2.16.1 featureCounts on curated simplified annotation format (SAF) interval files (promoter, gene-body, TSS-, TES-, TE-proximal, and consensus-peak intervals) (Liao *et al*., 2014). For TE intervals, reads were re-aligned in a multimapper-retaining mode (STAR, or bowtie2 --very-sensitive-local as fallback; MAPQ 0) and counted with featureCounts (countMultiMappingReads = TRUE, fraction = TRUE); all other intervals used uniquely mapped reads. Libraries isolated from their assigned stage or with the lowest within-stage Spearman correlation of 50-kb binned signal were excluded (two WT, one *ld-1*; Fig. S2a,b).

For differential accessibility, 1,470 candidate invariant genes were nominated from RPGC-normalized pooled promoter signal (absolute log2 fold-change < 0.25 and mean promoter RPGC ≥ 1.0 across WT torpedo-to-mature-green transitions; heart excluded), and DESeq2 size factors were estimated from the raw counts of these genes by geometric-mean ratios; VST values were used for PCA, clustering, and heatmaps. Metaprofiles around TSSs, TESs, gene bodies, and TE intervals were computed from RPGC-normalized BigWig files with rtracklayer (200 bins per flank and per scaled gene body; 10-bp resolution).

Accessibility modules were defined as for the transcriptome (stage-mean VST, feature-wise z-scores; k-means k = 4, 6, 8; set.seed(42), nstart = 50; ordered by peak stage).

A per-gene border-definition index (BDI) was computed from per-sample RPGC-normalized BigWig files. For each gene > 500 bp, BDI was the ratio of mean border signal (promoter −500/+200 bp of TSS and TES ±250 bp) to mean central gene-body signal (middle 60%), extracted with rtracklayer. Per-gene BDI was averaged across biological replicates within each stage and genotype (heart excluded from testing). Group differences were assessed by Kruskal–Wallis test (epsilon-squared effect size) with pairwise Wilcoxon rank-sum tests (Benjamini–Hochberg) and rank-biserial correlation (r) as a pairwise effect size.

Differential accessibility was tested using annotation-, window-, and union-peak-based approaches. Annotation-based count matrices were analysed with DESeq2 (genotype + stage designs). Genome-wide window-based testing used csaw v1.36.1 (150-bp sliding windows, 50-bp spacing, 5-kb background bins; readParam(pe = "both", dedup = TRUE, minq = 20)) with edgeR quasi-likelihood models (Robinson *et al*., 2010; Lun & Smyth, 2015). The bent-cotyledon group was reduced to two replicates after outlier exclusion, and T3 estimates are therefore conservative. For union-peak testing, per-stage reproducible peaks (MACS3 broad, overlapping ≥2 replicate peaks by bedtools multiinter) were summit-centred, extended to 500 bp, and merged into a non-redundant union set; counts (Rsubread featureCounts, paired-end, no multi-mapping) were tested with DESeq2 using the same control-gene size-factor estimation as the annotation-based approach.

Motif analyses were performed in two layers. Transcriptome-derived module and DE gene sets defined RNA-associated motif programs over promoter regions using the same monaLisa enrichment and TF-family normalization described above. Accessibility-side motif context used chromVAR bias-corrected deviation z-scores (Schep *et al*., 2017) and motif-centred accessibility quantification (below), with the same Display_Bin family definitions applied across layers. For genotype comparison, chromVAR z-scores were quantile-normalized per stage (limma::normalizeBetweenArrays) and tested with limma eBayes; the WT vs *ld-1* stage-mean z-score correlation (ρ) quantified preservation of the motif pattern. Accessible chromatin in the compact *Arabidopsis* genome concentrates at promoters (Sijacic *et al*., 2018; Bubb & Deal, 2020); annotation-based promoter windows (700 bp fixed) were therefore used as chromVAR input rather than called peaks.

### DNA methylation (WGBS) re-analysis

Public WGBS reads (Lee *et al*., 2023) were trimmed with Trim Galore (default --quality 20 --length 20) (Martin, 2011) and aligned to a bisulfite-converted *Arabidopsis* genome with Bismark v0.25.1 (bowtie2 backend) (Krueger & Andrews, 2011). Duplicates were removed with deduplicate_bismark, and per-context (CpG, CHG, CHH) methylation was called with bismark_methylation_extractor followed by bismark2bedGraph (--CX for CHG and CHH). A minimum coverage of 5 reads per cytosine was required before region- or tile-level summaries.

### Integrative analysis

Stage-matched transcriptome and NicE-seq data were integrated over a shared TAIR10/Araport11 framework, matching transcript abundance with promoter accessibility (500 bp upstream to 200 bp downstream of the TSS) and comparing stage-wise trajectories as normalized values or row-scaled z-scores.

Directed dependence between promoter accessibility and transcript abundance was quantified with the copula-based measure qad v1.0.4 (Junker *et al*., 2021), which estimates scale-invariant conditional predictability without assuming a functional form. Per-stage stage-mean values were supplied to qad::qad() at default resolution (excluding stages with < 50 genes); the directed values q(CA,Expr) and q(Expr,CA) and their difference provide a formal asymmetry measure. As a complementary metric, five-fold cross-validated isotonic regression (stats::isoreg; set.seed(42)) was performed in each direction, with R² computed on held-out folds. Reciprocal stratification ranked genes by promoter accessibility or expression within each stage and examined the other layer.

For cross-layer module concordance, layer-specific VST values were stage-averaged and z-scored over the core stages (heart excluded). Genes were grouped either by WT expression modules or by CA modules, and both expression and accessibility z-score trajectories were overlaid to assess whether grouping in one layer produced coherent trajectories in the other.

For motif-centred integration, motif sites within promoters were identified by scanning JASPAR 2024 plant PWMs with motifmatchr::matchMotifs (out = "positions", p.cutoff = 5e-5) on BSgenome.Athaliana.TAIR.TAIR9 (Rauluseviciute *et al*., 2024). For each module × TF-family combination, RPGC-normalized signal was extracted within ±75 bp of predicted motif sites, overlapping windows from the same family merged with GenomicRanges::reduce, and stage-specific z-scores computed against the genome-wide promoter mean and standard deviation for that family. Only module × TF-family combinations significantly enriched on the transcriptome side (adjusted P < 0.01) were included.

## Results

### Generation of stage-resolved transcriptome and CA datasets during mid-to-late *Arabidopsis* embryogenesis

We generated transcriptomic and genome-wide CA profiles from stage-resolved *Arabidopsis* embryos spanning heart, torpedo, bent-cotyledon, and mature-green stages, with developmental stages assigned according to standard morphological criteria (Fig. 1a). Transcriptome profiling was performed using Smart-seq3 on low-input RNA, whereas CA profiling used one-pot NicE-seq on fixed embryos. To evaluate data quality and stage-dependent relationships among samples, we performed principal component analysis (PCA) and computed pairwise correlations for both datasets. After quality filtering, transcriptomic profiles showed strong replicate concordance, and CA profiles exhibited clear stage-dependent separation (Fig. 1b–e).

**Fig. 1.**
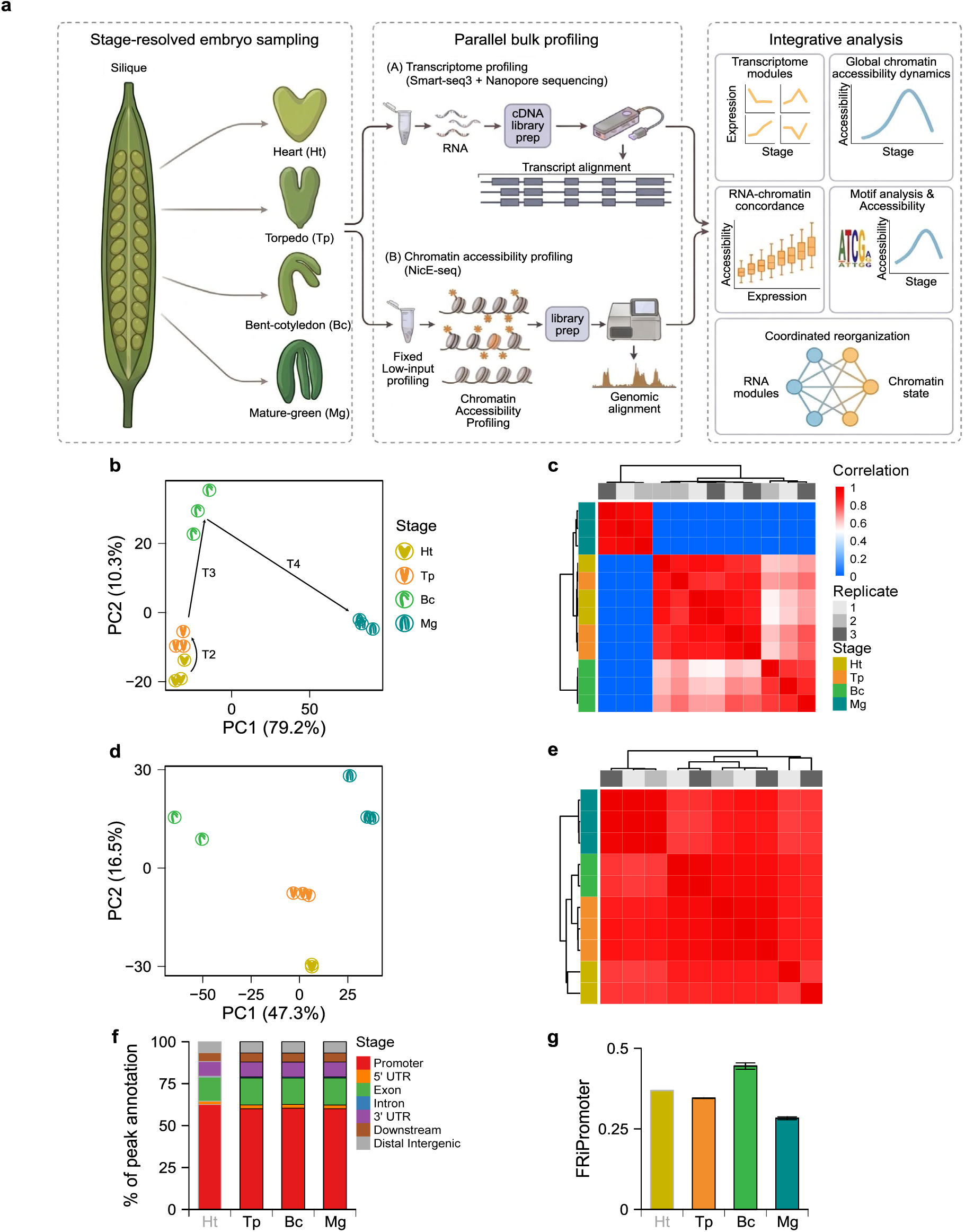
Experimental design, embryo staging, and dataset quality. **(a)** Schematic overview of the study design and sampled embryo stages: heart (Ht), torpedo (Tp), bent-cotyledon (Bc), and mature-green (Mg). **(b)** PCA of RNA-seq libraries based on the 2,000 most variable genes (VST values) across all biological replicates (n = 3 per stage). **(c)** Pairwise Spearman correlation heatmap of RNA-seq libraries computed from VST values. **(d)** PCA of NicE-seq libraries based on 50-kb binned chromatin accessibility signal across retained libraries (n = 3 per stage, except Ht and Bc, where one replicate was excluded after quality filtering). **(e)** Pairwise Spearman correlation heatmap of retained NicE-seq libraries computed from 50-kb binned accessibility signal. **(f)** Stage-resolved genomic annotation of MACS3 broad peaks showing the distribution of accessible regions across promoters, gene bodies, and intergenic regions. **(g)** FRiPromoter values across developmental stages, defined as the fraction of reads mapping to promoter regions from 500 bp upstream to 200 bp downstream of TSSs.

Transcriptomic variation was primarily structured by developmental stages, with PC1 and PC2 explaining 79.2% and 10.3% of the variance, respectively. The largest transcriptome-wide shift occurred during the T4 transition, followed by a substantial shift during T3, whereas the transcriptome-wide shift during T2 was comparatively limited (Fig. 1b,c).

CA profiles showed low within-stage concordance for one heart-stage replicate and one bent-cotyledon-stage replicate, which were therefore excluded from subsequent CA analyses. The remaining profiles showed clear stage-dependent clustering in PCA and correlation analyses (Fig. 1d,e; Supporting Information Fig. S2a,b; PC1 = 47.3%, PC2 = 16.5%). Torpedo, bent-cotyledon, and mature-green samples clustered reproducibly and were well resolved, whereas retained heart-stage replicates showed lower within-stage concordance and greater dispersion between replicates (Fig. S2a,b), consistent with lower signal recovery from early embryos. Accordingly, heart-stage libraries were retained only as an exploratory reference, and subsequent CA analyses focused on the later stages.

Annotation of accessible regions revealed strong enrichment near promoters and other gene-proximal regions (Fig. 1f; Fig. S3), consistent with previous reports that accessible chromatin in *Arabidopsis* is predominantly promoter-centric (Sijacic *et al*., 2018; Bubb & Deal, 2020). Accordingly, subsequent analyses quantified CA using annotation-defined promoter and gene-associated intervals rather than individual peak calls. FRiPromoter values (the fraction of reads falling within promoter regions, −500 to +200 bp relative to the transcription start site) peaked at the bent-cotyledon stage (∼44%) and declined at the mature-green stage (∼28%) (Fig. 1g). Together, these results indicate that CA remained strongly enriched at promoter-proximal regions throughout embryogenesis while undergoing substantial quantitative redistribution across developmental stages.

### Stage-specific transcriptome programs define mid-to-late embryogenesis

To characterize stage-resolved transcriptome dynamics during mid-to-late embryogenesis, we clustered genes into co-expression modules based on their expression trajectories across embryonic stages. Genes were grouped into eight co-expression modules (TX-M1 to TX-M8) to balance analytical resolution and biological interpretability while maintaining robust cluster structure (Fig. S1). These modules exhibited distinct temporal trajectories associated with different embryonic stages, indicating that the transcriptome during mid-to-late embryogenesis is organized into discrete developmental programs rather than a gradual continuum (Fig. 2a).

**Fig. 2.**
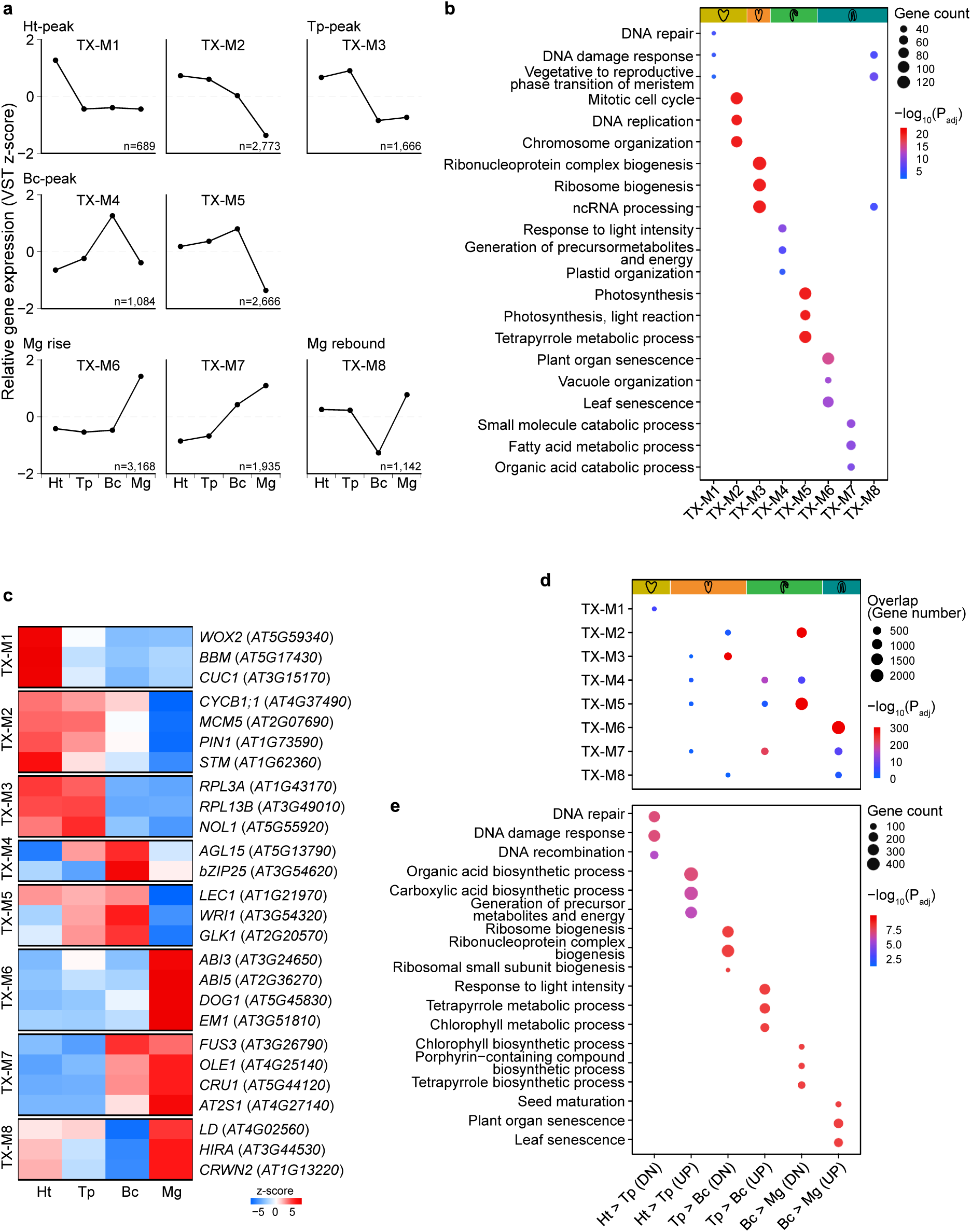
The transcriptome resolves *Arabidopsis* embryogenesis into stage-resolved developmental programs. **(a)** Expression module trajectories. Genes were grouped into eight modules by k-means clustering of stage-mean VST values row-scaled by gene (n = 3 biological replicates per stage). **(b)** GO Biological Process enrichment of the eight expression modules (clusterProfiler enrichGO, adjusted P < 0.05). Dot size indicates gene count, and color intensity indicates −log_10_(adjusted P). Top enriched terms for each module are shown. **(c)** Heatmap of selected representative genes and known regulatory anchors across all expression modules, highlighting representative developmental outputs, module identities, and major regulatory states. **(d)** Overlap between expression modules and stage-transition DE gene sets. For each consecutive transition (T2, T3, T4), upregulated and downregulated DE gene sets (DESeq2, FDR < 0.05) were tested for overlap with each module using one-sided Fisher’s exact test. Dot size indicates the number of overlapping genes, and color intensity indicates −log_10_(adjusted P). Significant enrichments (adjusted P < 0.05) are outlined. **(e)** GO Biological Process enrichment of stage-transition DE genes by GSEA. For each consecutive transition (T2, T3, T4), genes were ranked by the DESeq2 Wald statistic and tested against GO Biological Process terms. Dot size indicates gene count, and color intensity indicates −log10(adjusted P). Top enriched terms (adjusted P < 0.05) for each direction are shown.

Functional enrichment analysis linked these temporal modules to distinct developmental functions (Fig. 2b; Supporting Information Table S2). Early declining modules (TX-M1 and TX-M2) were enriched for DNA replication, embryo identity, cell cycle, and patterning functions, consistent with proliferative embryonic growth programs. A transient torpedo-peak module (TX-M3) was enriched for rRNA biogenesis and ribosome assembly, marking a metabolic transition between proliferative and maturation phases. Modules peaking at the bent-cotyledon stage (TX-M4 and TX-M5) were enriched for metabolic and plastid-related processes, consistent with the onset of embryo maturation, whereas late modules (TX-M6 and TX-M7) corresponded to reserve accumulation and dormancy-associated programs. In contrast, TX-M8 displayed a distinct biphasic trajectory, decreasing at the bent-cotyledon stage and rebounding at the mature-green stage, and was enriched for DNA damage response and chromatin organization rather than canonical reserve accumulation programs (Fig. 2a,b). Consistent with these functional assignments, selected representative genes and known developmental regulators showed module-specific expression timing that matched the corresponding developmental programs (Fig. 2c).

These temporal and functional patterns positioned TX-M4 and TX-M5 at the onset of maturation-associated expression programs, bridging earlier morphogenesis-associated modules and the late maturation modules that followed. This module boundary therefore captures the transcriptomic transition from morphogenesis-associated growth toward maturation. Consistent with these trajectories, transition-specific DE genes were strongly overrepresented in the corresponding temporal co-expression modules (Fig. 2d). Gene set enrichment analysis (GSEA) of transition-specific differentially expressed gene (DEG) sets further reinforced these functional assignments, linking early transitions to DNA replication and patterning, maturation onset to plastid and metabolic functions, and late embryogenesis to reserve accumulation and dormancy (Fig. 2e; Table S4).

Together, these findings indicate that mid-to-late embryogenesis is organized into discrete transcriptome programs, with a pronounced transition marking the shift from morphogenesis to maturation (Hofmann *et al*., 2019).

### Gene-associated accessibility undergoes global reorganization distinct from stage-specific transcriptome programs during embryo maturation

Having established that the transcriptome is organized into discrete stage-specific programs, we next asked whether the CA landscape mirrors this organization or follows a distinct trajectory. Examination of CA profiles across genic regions revealed a promoter-centered architecture with additional enrichment near polyadenylation sites (PASs). Such promoter- and PAS-proximal peaks progressively intensified toward the bent-cotyledon stage before collapsing at the mature-green stage (Fig. 3a).

**Fig. 3.**
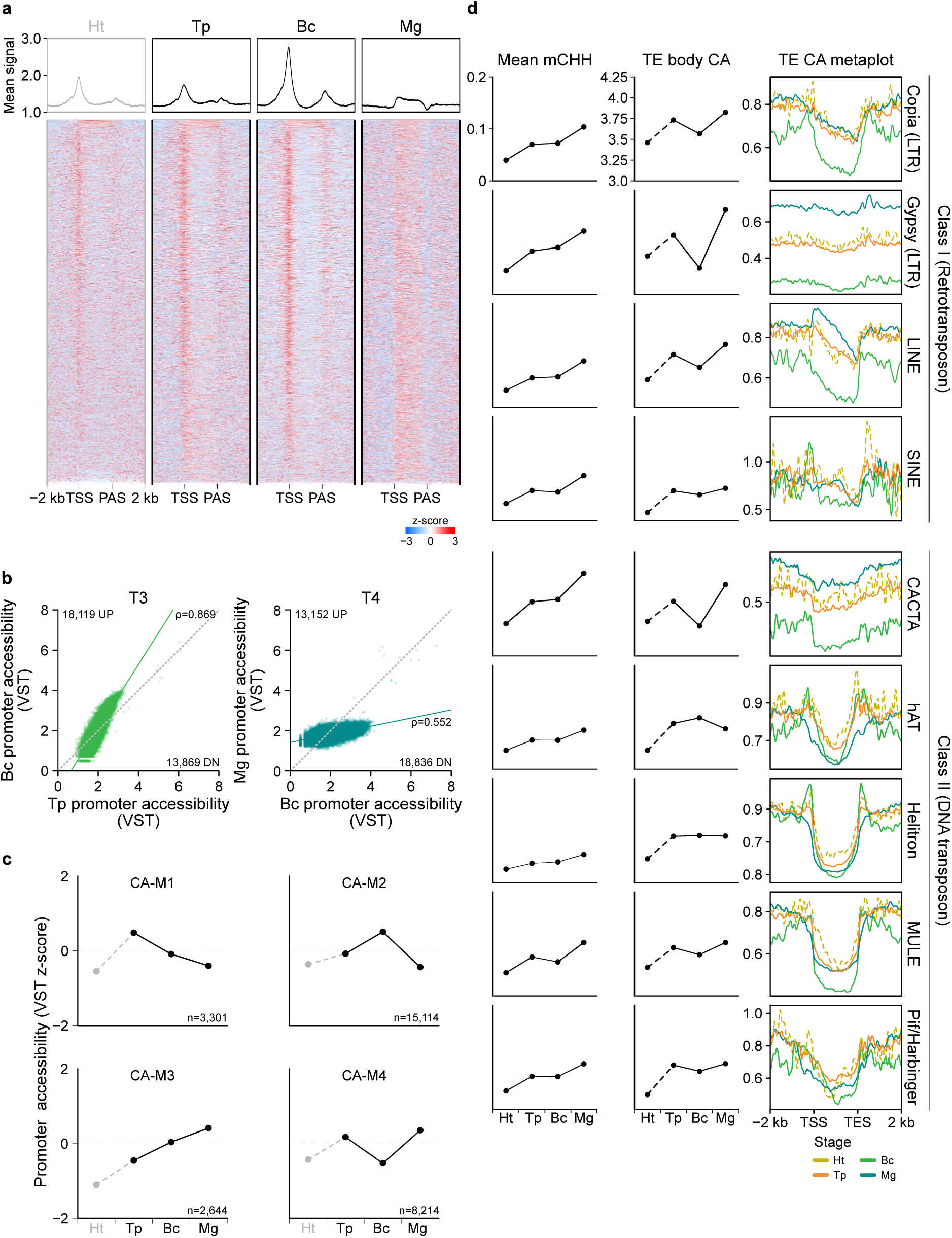
Gene-associated and TE-body accessibility undergo opposite dynamics during embryo maturation. **(a)** NicE-seq metaprofiles (RPGC-normalized signal) and gene-level heatmaps across developmental stages. Gene bodies were scaled to 2 kb with 2-kb flanking regions (200 bins per region; 10-bp effective resolution). Ht is shown as an exploratory reference, whereas Tp, Bc, and Mg represent the core stages used for downstream analyses. **(b)** Scatter plots of per-gene promoter accessibility (VST values from 500 bp upstream to 200 bp downstream of TSSs) between consecutive developmental stages. Each point represents one gene, and the dashed line indicates identity. Spearman correlation coefficients, total gene count (n = 31,988), and numbers of genes gaining or losing accessibility are annotated. **(c)** Accessibility module trajectory derived from k-means clustering of promoter-window accessibility z-scores across Tp, Bc, and Mg. Ht values are projected as an exploratory reference but were not used for module assignment. **(d)** CHH methylation dynamics from re-analysis of public WGBS datasets (Lee *et al*., 2023), TE-body chromatin accessibility trajectories, and accessibility metaprofiles across developmental stages for each TE family.

We then compared these accessibility profiles across stages to determine how they changed during embryo maturation. Because heart-stage libraries showed relatively low complexity, the apparent TSS (transcription start site) enrichment at this stage was interpreted cautiously and was not directly compared quantitatively with later stages (Fig. S2c,d). From the torpedo to bent-cotyledon stages, CA profiles progressively sharpened: TSS peak height increased by ∼1.5-fold, PAS-proximal enrichment became clearly defined, and the contrast between promoter-proximal and gene-body signal peaked at the bent-cotyledon stage. Gene-level heatmaps indicated that this sharpening was widespread across the genome rather than restricted to a small subset of highly accessible loci (Fig. 3a). In contrast, mature-green embryos deviated from this trajectory and showed collapse of the promoter-centered accessibility pattern. TSS- and PAS-proximal enrichment was largely lost, gene-body signal rose slightly relative to flanking regions, and the overall promoter-centered accessibility profile became flattened (Fig. 3a). Together, these results indicate that promoter- and PAS-centered CA organization sharpened from torpedo to bent-cotyledon before collapsing at the mature-green stage.

Because the mature-green stage displayed an atypical promoter accessibility profile, we next asked whether this apparent collapse could reflect a technical artifact. However, mature-green libraries exhibited the highest complexity among all stages, arguing against low library complexity as the cause of this pattern (Fig. S2c). To further exclude read-depth effects, all libraries were downsampled to the unique read count of the lowest-depth core-stage library (bent-cotyledon; 1.17M unique reads). This analysis reproduced the same stage-dependent accessibility trajectory observed in the original dataset (Fig. S2f). Subsequent CA analyses were therefore performed using the original non-downsampled libraries from the torpedo, bent-cotyledon, and mature-green stages.

To determine whether the global metaprofile changes were recapitulated at the level of individual genes, we quantified promoter accessibility across embryonic stage transitions (Fig. 3b; Fig. S2e for the exploratory T2 transition). Gene-level analyses confirmed that the accessibility changes observed in the global metaprofiles reflected a genome-wide redistribution of accessibility rather than changes restricted to a limited subset of loci. At T3, ∼57% of genes showed increased promoter accessibility (mean gain: 0.57 variance-stabilized transformation [VST] units), whereas at T4 this pattern reversed, with ∼59% of genes showing decreased accessibility (mean loss: 0.89 VST units), despite substantial bidirectional accessibility changes at both transitions.

In contrast to the stage-specific transcriptome programs, promoter accessibility dynamics were largely governed by a shared global trajectory. To examine this structure in greater detail, we grouped genes into four accessibility modules (CA-M1 to CA-M4) based on their promoter accessibility trajectories across the core stages (Fig. 3c; Fig. S2g; Table S5). More than half of genes (∼52%; 15,114 genes) belonged to a single accessibility module, CA-M2, which followed the dominant pattern of a bent-cotyledon peak followed by a sharp decline at the mature-green stage. The remaining modules followed related but distinct trajectories. CA-M1 peaked at the torpedo stage and declined thereafter, CA-M3 progressively increased toward the mature-green stage, and CA-M4 showed reduced accessibility at the bent-cotyledon stage followed by partial recovery at the mature-green stage.

Unlike the transcriptome modules, Gene Ontology (GO) enrichment of the accessibility modules did not partition into similarly distinct stage-specific developmental programs (Fig. S4; Table S6). Instead, the observed accessibility changes appear to reflect a broadly shared structural transition rather than distinct chromatin programs associated with separate developmental functions.

Taken together, these results indicate that *Arabidopsis* embryo maturation involves a two-phase reorganization of gene-associated accessibility: promoter- and PAS-proximal accessibility profiles sharpen from torpedo to bent-cotyledon, reaching maximal promoter–gene-body contrast before collapsing during the transition toward seed dormancy.

### Transposable element body accessibility exhibits dynamics broadly opposite to promoter accessibility

We next examined accessibility dynamics of transposable elements (TEs) during mid-to-late *Arabidopsis* embryogenesis. TE regulation during plant embryogenesis has been studied primarily at the level of DNA methylation, particularly the progressive accumulation of CHH methylation associated with maturation-linked silencing (Bouyer *et al*., 2017; Kawakatsu *et al*., 2017). We therefore asked whether TE body accessibility progressively declines during embryo maturation as CHH methylation accumulates or instead follows distinct developmental dynamics.

To compare TE accessibility dynamics with maturation-associated DNA methylation changes, we re-analyzed public whole-genome bisulfite sequencing (WGBS) data from matched embryonic stages. This analysis confirmed previously reported TE methylation patterns: CG and CHG methylation remained relatively stable, whereas CHH methylation gradually increased (Fig. 3d; Fig. S5a) (Lee *et al*., 2023). In contrast to this methylation trend, TE body accessibility did not progressively decline but instead followed distinct family-specific trajectories across embryonic stages. Because these accessibility changes could potentially reflect altered TE transcriptional activity, we also considered TE expression dynamics. However, after exclusion of gene-overlapping reads, the number of confidently assigned autonomous TE-derived transcripts was too low for robust locus-level quantification (Fig. S5b). We therefore interpreted TE dynamics primarily at the level of chromatin accessibility rather than attempting locus-level inference of TE transcriptional activity.

To compare TE dynamics with the promoter accessibility trajectory described above, TE body accessibility was examined across developmental stages. In most TE families, stage-resolved TE body accessibility followed trajectories broadly opposite to promoter accessibility dynamics (Fig. 3d). Whereas promoter accessibility peaked at the bent-cotyledon stage and sharply declined at the mature-green stage (Fig. 3a), most TE families, including Gypsy, Copia, LINE, SINE, CACTA, MULE, and Pif/Harbinger, showed the lowest accessibility at the bent-cotyledon stage, followed by increased accessibility at the mature-green stage. In contrast, hAT and Helitron elements deviated from this pattern: hAT showed gene-like boundary-proximal accessibility, whereas Helitron displayed a relatively stable trajectory (Fig. 3d).

Together, these results indicate that TE body accessibility follows family-specific dynamics broadly opposite to gene-associated promoter accessibility. These patterns cannot be explained by CHH methylation accumulation alone and instead suggest distinct modes of accessibility remodeling in gene-associated regions and TEs during embryo maturation.

### Promoter accessibility largely follows a global trajectory, with limited motif-level correspondence to expression

Having established that the transcriptome is organized into stage-specific programs whereas accessibility at gene promoters follows a largely global trajectory, we next examined how strongly promoter accessibility and transcription are coupled during embryo maturation and whether this relationship becomes more apparent at the level of transcription factor (TF)-binding motifs (Fig. 4).

**Fig. 4.**
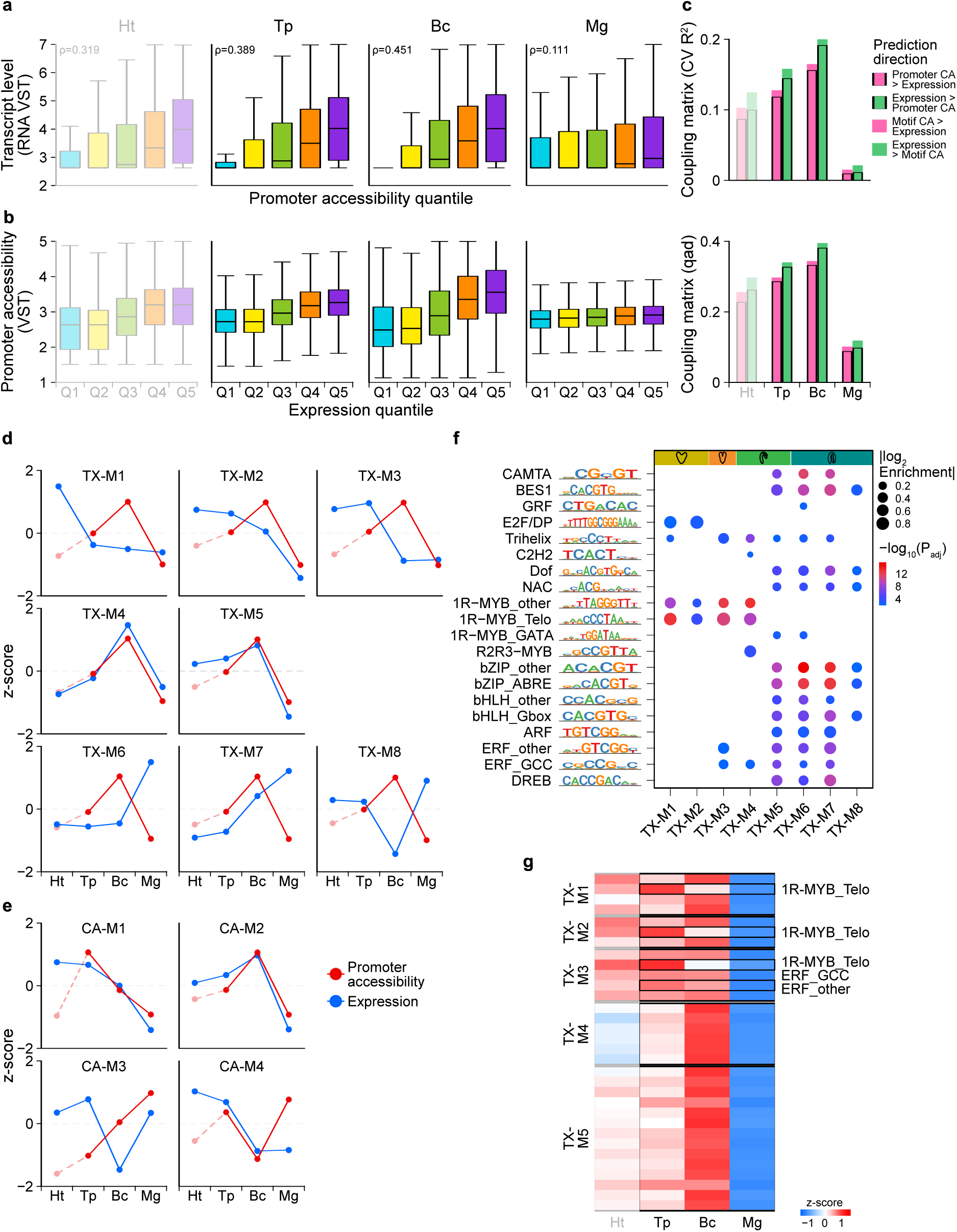
Promoter and motif accessibility follow a global trajectory with limited module-specific correspondence. **(a)** Genes were ranked by promoter accessibility quintile (VST) within each developmental stage, and transcript abundance for each quintile is shown as boxplots (center line, median; box, IQR; whiskers, 1.5× IQR). Spearman’s rank correlation (ρ) between promoter accessibility and transcript level is annotated for each stage. Stage-wise mean values across biological replicates are shown (n = 3 per stage, except Bc NicE-seq, n = 2 after quality filtering). **(b)** Genes were ranked by expression quintile (RNA VST) within each developmental stage, and promoter accessibility for each quintile is shown as boxplots (center line, median; box, IQR; whiskers, 1.5× IQR). **(c)** Per-gene promoter-level and per-motif accessibility–transcription coupling across developmental stages, quantified using qad and five-fold cross-validated isotonic regression (CV R²). Promoter-level coupling was calculated using promoter accessibility (500 bp upstream to 200 bp downstream of TSSs), whereas motif-level coupling used local accessibility within ±75 bp of predicted motif sites paired with expression of the corresponding host genes. Both prediction directions are shown for each stage. Ht is included as an exploratory reference. **(d)** Module-level correspondence between accessibility and expression using expression-defined modules. Accessibility trajectories are shown for genes grouped by expression module. Gene expression is shown in blue and promoter accessibility in red; values are displayed as z-scores across developmental stages. **(e)** Module-level correspondence between accessibility and expression using accessibility-defined modules. Expression trajectories are shown for genes grouped by accessibility module. Gene expression is shown in blue and promoter accessibility in red; values are displayed as z-scores across developmental stages. **(f)** Promoter motif enrichment across expression modules based on RNA-defined gene sets (JASPAR 2024, motif-subfamily collapse, adjusted P < 0.01). Dot size indicates enrichment score, and color indicates −log10(adjusted P). **(g)** Module × TF-family promoter accessibility z-scores across Tp, Bc, and Mg relative to the genome-wide stage mean. The 32 rows correspond to the TX-M1–TX-M5 subset of the 66 module × TF-family pairs that passed motif enrichment significance (adjusted P < 0.01 in Fig. 4f). TX-M6–TX-M8 are shown in Fig. S6.

At the individual gene level, genes with higher promoter accessibility generally showed higher transcript abundance across all developmental stages (Fig. 4a), although the strength of this relationship varied substantially during embryo maturation. Concordance between promoter accessibility and transcript abundance was strongest at the bent-cotyledon stage (ΔQ5–Q1 = 1.35 variance-stabilized transformation [VST] units) and weakest at the mature-green stage (0.35 VST units). Spearman correlation (ρ) showed a similar pattern, increasing from torpedo to bent-cotyledon before dropping sharply at the mature-green stage (ρ = 0.39, 0.45, and 0.11, respectively; Fig. 4a). Reciprocal analysis further showed that promoter accessibility differences were most apparent among highly expressed genes, whereas genes in the lower expression quintiles (Q1–Q2) showed relatively little difference in promoter accessibility compared to highly expressed quintiles (Q3–Q5; Fig. 4b). Quantification using qad (asymmetric dependence) and five-fold cross-validated isotonic regression (CV R²; predictive strength) confirmed stage-dependent CA–transcription coupling, with both metrics peaking at bent-cotyledon before declining to minimal levels at mature-green (qad ≤ 0.10; CV R² ≤ 0.011; Fig. 4c).

We next asked whether promoter accessibility reflects the stage-specific transcriptome programs identified above. When genes were grouped according to transcriptome modules (TX-M1–TX-M8), all modules showed highly similar promoter accessibility trajectories despite their distinct expression patterns (Fig. 4d). Conversely, grouping genes according to accessibility modules (CA-M1–CA-M4) failed to resolve distinct expression programs (Fig. 4e). These reciprocal analyses indicate that promoter accessibility and transcription remain only partially coupled at the module level.

To determine whether stronger concordance emerges at finer regulatory resolution, we performed motif enrichment analysis within each transcriptome module (Fig. 4f; Table S7). Early modules (TX-M1–TX-M4) were enriched primarily for 1R-MYB motifs, with E2F/DP motifs specifically enriched in TX-M1 and TX-M2, consistent with E2F–RBR-mediated cell-cycle exit during embryogenesis (Leviczky *et al*., 2019). Motif composition shifted between TX-M4 and TX-M5, after which 1R-MYB signatures were replaced by motifs associated with seed maturation regulators, including bZIP, BES1, bHLH, DREB, ERF, CAMTA, Dof, NAC, and ARF families (Holdsworth *et al*., 2008; Santos-Mendoza *et al*., 2008; Alonso *et al*., 2009). Thus, transcriptome modules were associated with distinct motif-level regulatory signatures.

To test whether these motif-defined regulatory signatures were reflected in the chromatin landscape, we quantified promoter accessibility within ±75 bp of each predicted motif site and compared each module–TF combination against the stage-specific genome-wide mean accessibility for the corresponding motif family (Fig. 4g). Despite the distinct motif enrichments observed across transcriptome modules, 61 of 66 module–motif combinations showed maximal accessibility at the bent-cotyledon stage (Fig. 4g; Fig. S6). Only five combinations deviated from this pattern, all within early torpedo-associated modules (TX-M1–TX-M3), where motif accessibility aligned more closely with the expression timing of the corresponding host module than with the global accessibility trajectory. These exceptions indicate that motif-specific accessibility can locally override the shared accessibility trajectory.

Together, these results indicate that promoter accessibility and transcription are only partially coupled during *Arabidopsis* embryo maturation. Although motif-level analysis revealed discrete regulatory signatures, most accessibility patterns remained dominated by a shared global accessibility trajectory, with only a limited subset of motif-associated accessibility profiles locally aligning with module-specific expression timing.

### Loss of *LD* reconfigures motif accessibility toward expression-concordant timing

Our WT analyses showed that chromatin accessibility (CA) and transcription are only partially coupled during embryo maturation. To test whether this limited concordance reflects a fixed chromatin property or a developmentally regulated state, we examined the *ld-1* mutant of the global transcriptional repressor *LUMINIDEPENDENS*, a gene associated with transcriptional and chromatin regulatory pathways (He *et al*., 2003; Liu *et al*., 2007; Mateo-Bonmatí *et al*., 2024; Bergis-Ser *et al*., 2025). This allowed us to test whether the strength of CA–transcription coupling can be genetically reconfigured in embryonic chromatin.

We first examined *LD* expression across embryonic stages. *LD* expression was lowest at the bent-cotyledon stage, when promoter accessibility was maximal, and rebounded at the mature-green stage, when accessibility declined (Fig. 5a). PCA showed that WT and *ld-1* transcriptomes largely followed the same developmental trajectory, although *ld-1* bent-cotyledon samples shifted slightly toward the torpedo-stage cluster (Fig. 5b). Consistent with this overall preservation of developmental structure, only modest transcriptional changes were detected across stages, with 38, 36, 119, and 163 DEGs identified at the heart, torpedo, bent-cotyledon, and mature-green stages, respectively (Table S8). This modest count likely reflects the library-size normalization inherent to standard RNA-seq, which cannot capture genome-wide shifts in transcript abundance; spike-in–based profiling has shown that loss of LD causes widespread transcriptional derepression (Bergis-Ser et al., 2025). Our analyses therefore focus not on the magnitude of expression change but on its developmental timing and its coupling to accessibility, which are unaffected by this normalization.

**Fig. 5.**
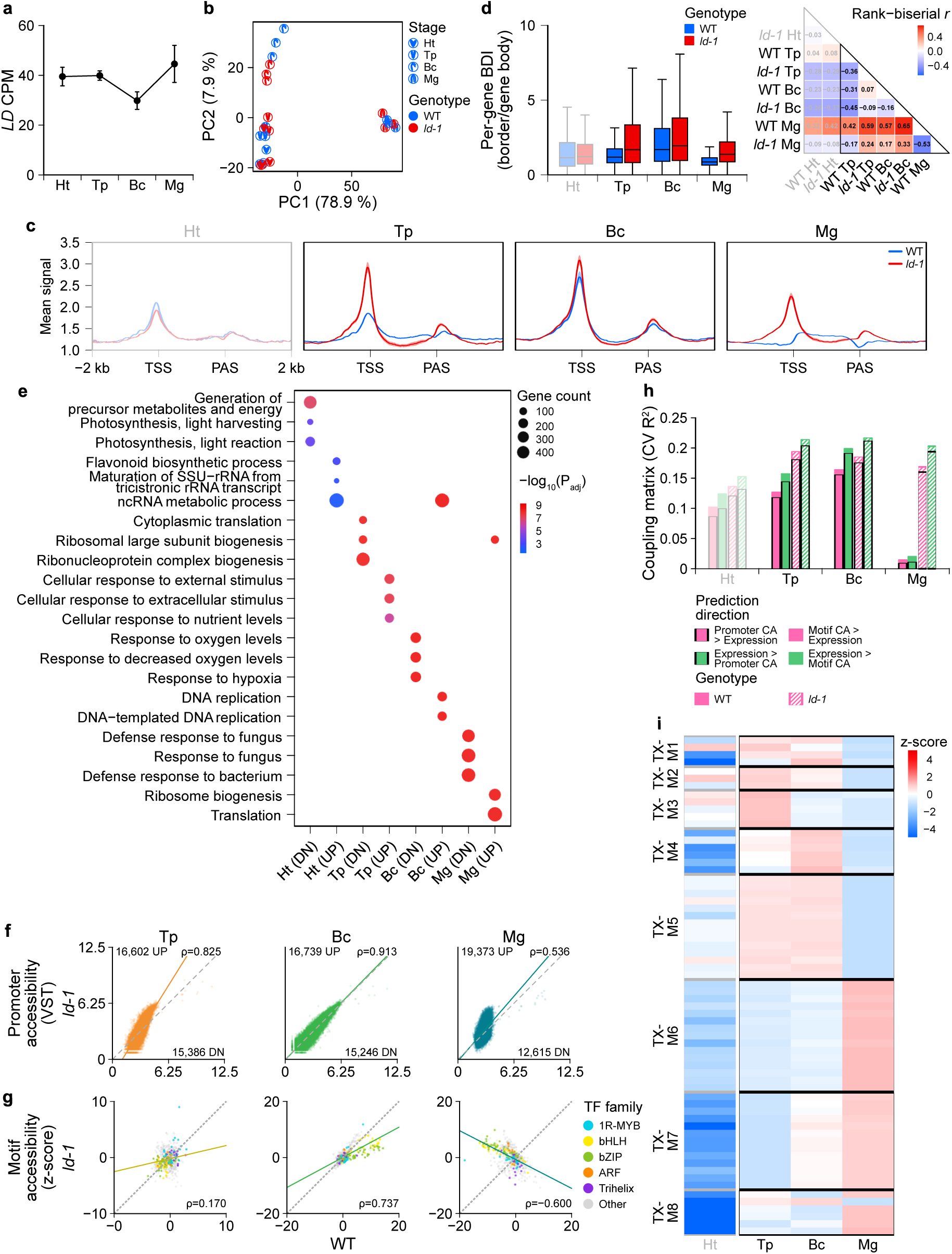
Loss of *LD* reconfigures motif accessibility toward expression-concordant timing. **(a)** *LD* (*AT4G02560*) expression trajectory across four embryonic stages in WT (mean CPM ± SE, n = 3). **(b)** PCA of the 2,000 most variable genes for WT and *ld-1* samples. **(c)** NicE-seq metaprofiles (RPGC-normalized signal) of WT and *ld-1* across developmental stages using the same binning strategy as in Fig. 3a. Each panel shows one developmental stage with both genotypes overlaid (WT, blue; *ld-1*, red). Ht is included as an exploratory reference. **(d)** Per-gene border-definition index (BDI) across developmental stages for WT and *ld-1* (left) and pairwise rank-biserial effect sizes (r) for all genotype × stage comparisons (right). BDI was calculated as the ratio of border signal to central gene-body signal (middle 60%), with higher values indicating sharper promoter- and PAS-proximal enrichment. Per-gene BDI values were averaged across biological replicates before statistical testing. Genotype differences within each stage were assessed using Wilcoxon rank-sum tests. **(e)** GSEA of genotype-dependent transcriptional differences at each developmental stage. Genes were ranked by the DESeq2 Wald statistic (*ld-1* versus WT) and tested against GO Biological Process terms. Top enriched terms (adjusted P < 0.05) are shown. **(f)** Scatter plots of per-gene promoter accessibility (VST) between WT and ld-1 across Tp, Bc, and Mg. Stage-mean VST values were computed for each genotype without quantile normalization. Each point represents one gene; dashed lines indicate identity, and solid lines indicate linear regression fits. Spearman correlation coefficients and numbers of genes above and below the diagonal are annotated. **(g)** Scatter plots of chromVAR motif deviation z-scores between WT and *ld-1* across Tp, Bc, and Mg. Each point represents one motif; dashed lines indicate identity, and solid lines indicate linear regression fits. Points are colored by TF family (1R-MYB, bHLH, bZIP, ARF, and Trihelix; remaining families shown in grey). Spearman correlation coefficients between WT and ld-1 stage-mean z-scores are annotated. **(h)** Comparison of per-gene promoter-level and per-motif accessibility–transcription coupling between WT and *ld-1* across developmental stages using the framework described in Fig. 4c. CV R² metrics and both prediction directions are shown for each genotype and stage. Ht is included as an exploratory reference. **(i)** Module × TF-family promoter accessibility z-scores in *ld-1* across Tp, Bc, and Mg using the same analysis framework as in Fig. 4g.

In contrast to these limited transcriptome changes, CA was substantially altered in *ld-1* (Fig. 5c; Fig. S7a,b). At the torpedo stage, *ld-1* already showed strong promoter-and PAS-proximal accessibility comparable to the WT bent-cotyledon peak. Unlike WT, which showed flattening of promoter accessibility profiles at mature-green, *ld-1* maintained strong TSS- and PAS-proximal enrichment. To quantify these changes at the single-gene level, we calculated a border-definition index (BDI), defined as the ratio of border signal (promoter and PAS ± 250 bp) to central gene-body signal (middle 60%) (Fig. 5d). In WT, BDI increased from torpedo to bent-cotyledon and declined below 1 at mature-green. In contrast, *ld-1* showed elevated BDI already at torpedo and maintained values above 1 at mature-green. Thus, loss of *LD* caused persistent strengthening of promoter- and PAS-proximal boundary structure across embryonic stages.

We next asked whether these accessibility changes were accompanied by coordinated transcriptional shifts. Although relatively few genes were differentially expressed, gene set enrichment analysis (GSEA) revealed altered developmental timing of transcriptional programs (Fig. 5e; Table S9). Ribosome biogenesis programs were prematurely downregulated at torpedo but remained enriched at mature-green in *ld-1* relative to WT. Nevertheless, canonical maturation markers converged between WT and *ld-1* by the mature-green stage (Fig. S7c), indicating that loss of *LD* alters transcriptional timing without disrupting the overall maturation program.

We next compared WT and *ld-1* accessibility patterns at promoter and motif levels. Promoter-level accessibility patterns remained broadly similar between WT and *ld-1*, consistent with strong stage-dependent positive correlations (ρ = 0.83, 0.91, and 0.54 at torpedo, bent-cotyledon, and mature-green, respectively; Fig. 5f). In contrast, motif-level accessibility patterns were substantially altered in *ld-1*, showing weakened or reversed stage-wise correlations relative to WT across developmental stages (ρ = 0.17, 0.74, and −0.60 at torpedo, bent-cotyledon, and mature-green, respectively; Fig. 5g; Table S10). Thus, motif-proximal accessibility was more sensitive to loss of *LD* than broader promoter accessibility.

Because WT-defined module trajectories remained interpretable in *ld-1* (Fig. S7d,e), we next examined whether CA–transcription coupling was altered in the mutant. Coupling was quantified using the same metrics as in Fig. 4c (Fig. 5h; Fig. S7f). In WT, promoter- and motif-level coupling peaked at bent-cotyledon and declined sharply at mature-green. In contrast, *ld-1* showed consistently stronger coupling across all stages, with the greatest increase at mature-green. Coupling also remained consistently stronger at the motif level than at the promoter level in both WT and *ld-1*.

Finally, we asked whether loss of *LD* alters the relationship between motif accessibility and module-specific expression timing. In WT, most motif accessibility profiles followed a shared global trajectory largely independent of transcriptome modules (Fig. 4g). In *ld-1*, however, many motif-associated accessibility profiles shifted toward the transcriptional timing of their host modules (Fig. 5i). Early modules (TX-M2 and TX-M3) shifted from the WT bent-cotyledon-centered accessibility pattern toward earlier torpedo-stage accessibility in *ld-1*, most prominently in TX-M3. Conversely, late modules (TX-M6–TX-M8) shifted toward maximal accessibility at mature-green, with 33 of 34 module–motif combinations showing mature-green accessibility peaks. Thus, loss of *LD* increased alignment between motif accessibility and module-specific expression timing.

Together, these results indicate that CA–transcription coupling is not a fixed property of embryonic chromatin but a regulated state that can be genetically reconfigured. Loss of *LD* enhances CA–transcription coupling, particularly at the motif level, and shifts accessibility dynamics toward module-specific expression timing.

## Discussion

In this study, we define CA–transcription coupling as a dynamically regulated property of *Arabidopsis* embryo maturation rather than as a fixed reflection of transcriptional activity. Stage-resolved profiling revealed that promoter- and PAS-proximal accessibility followed a globally coordinated trajectory that peaked at the bent-cotyledon stage and collapsed at the mature-green stage, whereas transcriptome organization resolved into discrete stage-specific developmental programs. Although most accessibility dynamics were dominated by this shared global trajectory, motif-level analyses identified a limited subset of accessibility patterns locally aligned with module-specific expression timing. Importantly, loss of *LD* further enhanced this alignment between accessibility and transcription. Together, these findings indicate that embryo maturation involves a globally coordinated accessibility state that is only partially coupled to transcription but remains developmentally and genetically reconfigurable.

This study represents an early application of NicE-seq to stage-resolved plant embryonic tissues. Using a simplified one-pot workflow without nuclei purification (Vishnu *et al*., 2021), we generated reproducible accessibility profiles from torpedo, bent-cotyledon, and mature-green embryos. Although heart-stage libraries showed lower signal recovery and replicate concordance, likely reflecting limited nuclear input, the overall approach proved compatible with fixed low-input embryonic material. Because intact nuclei isolation is often technically challenging in plant developmental tissues, this workflow may facilitate future accessibility profiling in similarly limited or difficult-to-isolate samples.

A major finding of this study is that gene-associated accessibility during embryo maturation follows a globally coordinated accessibility state that is largely independent of module-specific transcriptional organization. Promoter- and PAS-proximal accessibility progressively sharpened from torpedo to bent-cotyledon before collapsing at mature-green, whereas transcriptome modules followed discrete developmental programs associated with morphogenesis, maturation onset, and dormancy preparation. More than half of all genes belonged to a single accessibility module characterized by this bent-cotyledon-centered accessibility pattern, and accessibility modules showed relatively limited partitioning into distinct biological functions. Our data therefore support a model in which embryo maturation involves a globally coordinated accessibility state with only limited correspondence to stage-specific transcriptional programs.

The disconnect between globally coordinated accessibility and module-specific transcriptional programs became particularly evident in motif-centered analyses.

Transcriptome modules carried distinct TF-family enrichments associated with morphogenesis and seed maturation programs. However, motif accessibility itself largely followed the same bent-cotyledon-centered global trajectory regardless of module identity. Thus, distinct TF-family enrichments were superimposed onto a largely shared accessibility landscape in WT embryos. This mismatch was especially pronounced in late maturation modules, where TF-family signatures associated with mature-green transcriptional programs nevertheless showed maximal motif accessibility at the bent-cotyledon stage. Together with the asymmetric predictive relationship observed between accessibility and expression, these findings suggest that accessibility during embryo maturation primarily establishes a permissive developmental context within which transcriptional specificity is subsequently imposed by TF activity and other regulatory inputs.

The partial coupling between CA and transcription may partly reflect the compact regulatory architecture of the *Arabidopsis* genome. In *Arabidopsis*, accessible chromatin is concentrated predominantly near promoters (Sijacic *et al*., 2018; Bubb & Deal, 2020), and our analyses further showed that accessibility trajectories were largely shared across transcriptome modules rather than tailored to individual developmental programs. Similar partial coupling between accessibility and transcription has also been reported in other *Arabidopsis* contexts, including shade responses and natural accession variation (Alexandre *et al*., 2018; Paulišić *et al*., 2025). Together, these observations support a model in which compact promoter-centric genomes such as *Arabidopsis* favor broadly permissive accessibility states with only limited coupling to module-specific transcriptional programs.

During embryo maturation, gene-associated and TE-associated accessibility underwent distinct but coordinated reconfiguration. Whereas gene-associated accessibility followed the globally coordinated trajectory centered on the bent-cotyledon stage described above, most TE families showed broadly opposite accessibility dynamics despite progressive CHH methylation during embryo maturation. These findings suggest that maturation-associated accessibility remodeling differs fundamentally between gene-associated regions and TE bodies. Nonetheless, because these analyses rely on normalized accessibility profiles, we cannot fully distinguish absolute changes in TE accessibility from compositional redistribution among genomic regions; apparent family-level trajectories may also partly reflect TE length or subclass composition rather than superfamily identity alone.

Placing these findings within plant embryogenesis more broadly, our *Arabidopsis* dataset shares core features with previous CA profiling during wheat embryogenesis (Zhao *et al*., 2023): in both systems, CA rose during mid-embryogenesis before undergoing pronounced restriction during late maturation. However, accessibility and transcription were more concordant in wheat than in *Arabidopsis*, which may reflect differences in genome architecture between compact gene-dense genomes and large genomes with abundant distal regulatory space (Alexandre *et al*., 2018; Sijacic *et al*., 2018; Ricci *et al*., 2019; Bubb & Deal, 2020; Zhao *et al*., 2023; Paulišić *et al*., 2025). While maturation-associated accessibility restriction may be a shared feature of plant embryogenesis, the degree of CA–transcription coupling appears to depend on genome architecture.

The maturation-associated accessibility collapse observed here also differs from many animal developmental systems. In animal embryogenesis and differentiation, CA is generally established progressively across regulatory elements and cell states, without a comparable maturation-associated global accessibility collapse (Klemm *et al*., 2019; Lynch *et al*., 2022; Pollex *et al*., 2024). Such differences may reflect distinct developmental modes: most animal organ systems are established during embryonic development, whereas plant organogenesis remains incomplete before post-embryonic growth and requires global chromatin stabilization. Nonetheless, partial CA–transcription coupling itself appears to be a recurring feature of developmental systems across kingdoms (Klemm *et al*., 2019; Lynch *et al*., 2022; Pollex *et al*., 2024).

Genetic perturbation in *ld-1* demonstrates that this limited CA–transcription coupling is not a fixed property of embryonic chromatin. In *ld-1*, promoter- and PAS-proximal accessibility became prematurely strengthened and remained despite only modest differential expression at the detection level. Indeed, because LD-dependent derepression is largely global and thus underrepresented by standard normalization, the accessibility layer is reconfigured even where differential expression is not registered, underscoring that the control of CA is separable from transcriptional output. More importantly, motif accessibility became more strongly aligned with module-specific expression timing, and overall CA–transcription concordance increased across developmental stages. These findings indicate that the permissive accessibility state observed in WT embryos can be genetically reconfigured toward a more expression-concordant state.

This reconfiguration also dissociates CA timing from maturation identity. Although canonical maturation markers converged by mature-green, promoter architecture and motif-level accessibility remained displaced, arguing against a purely passive model of accessibility collapse during seed maturation. This interpretation is consistent with recent studies showing that LD acts through multiple regulatory mechanisms, including RNA polymerase II regulation, Mediator-associated transcription, and FLD-mediated chromatin regulation (He *et al*., 2003; Liu *et al*., 2007; Mateo-Bonmatí *et al*., 2024; Bergis-Ser *et al*., 2025). Rather than acting solely as a local chromatin-closing factor, LD may contribute to a buffering layer that limits the extent to which TF-driven transcriptional states are reflected in motif-proximal accessibility.

Loss of this buffering capacity may also influence TF occupancy patterns by disproportionately opening otherwise weakly accessible loci, thereby redistributing the accessibility landscape experienced by TFs (Long *et al*., 2023; Taylor *et al*., 2024). Together, these observations support a hierarchical regulatory model in which CA provides developmental competence and regulatory potential, whereas transcriptional outputs are determined by the balance between activating and repressive TF inputs operating within that accessible state. In this view, the long-debated discordance between CA and transcription reflects a developmentally regulated property rather than a fixed feature of chromatin organization — one that can be genetically reconfigured by perturbing the global regulatory layers that buffer this relationship.

## Acknowledgements

We thank Junghoon Park and Dae-Jin Yun (Konkuk University) for advice on NicE-seq library preparation, and Yeonhee Choi and colleagues for making embryo WGBS data publicly available (Lee *et al*., 2023). This work was supported by the National Research Foundation (NRF) of Korea (RS-2021-NR060084 and RS-2025-00520643).

## Author contributions

DJ, CP, and IL conceived the study and designed the research. DJ performed the experiments, analyzed the data, and prepared the original manuscript draft. DJ, CP, and IL interpreted the results and revised the manuscript. CP and IL supervised the project. All authors read and approved the final manuscript.

## Data availability

Nanopore RNA-seq and NicE-seq data have been deposited in NCBI under BioProject PRJNA1470972. Public WGBS data were obtained from NCBI BioProject PRJNA944400 (Lee *et al*., 2023).

## Supplementary Figures

**Fig. S1.**
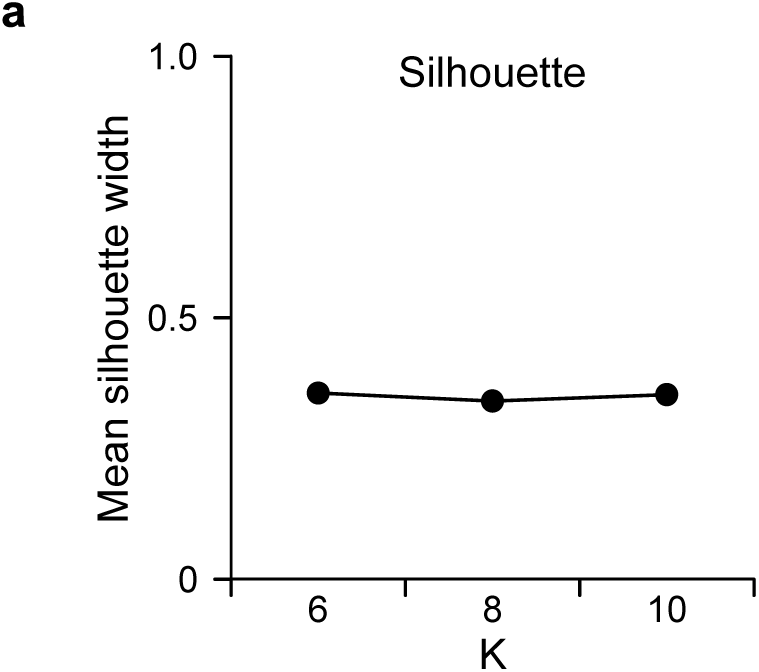
Expression module robustness. **a** Cluster-quality diagnostics for expression module clustering with k = 6, 8, and 10.

**Fig. S2.**
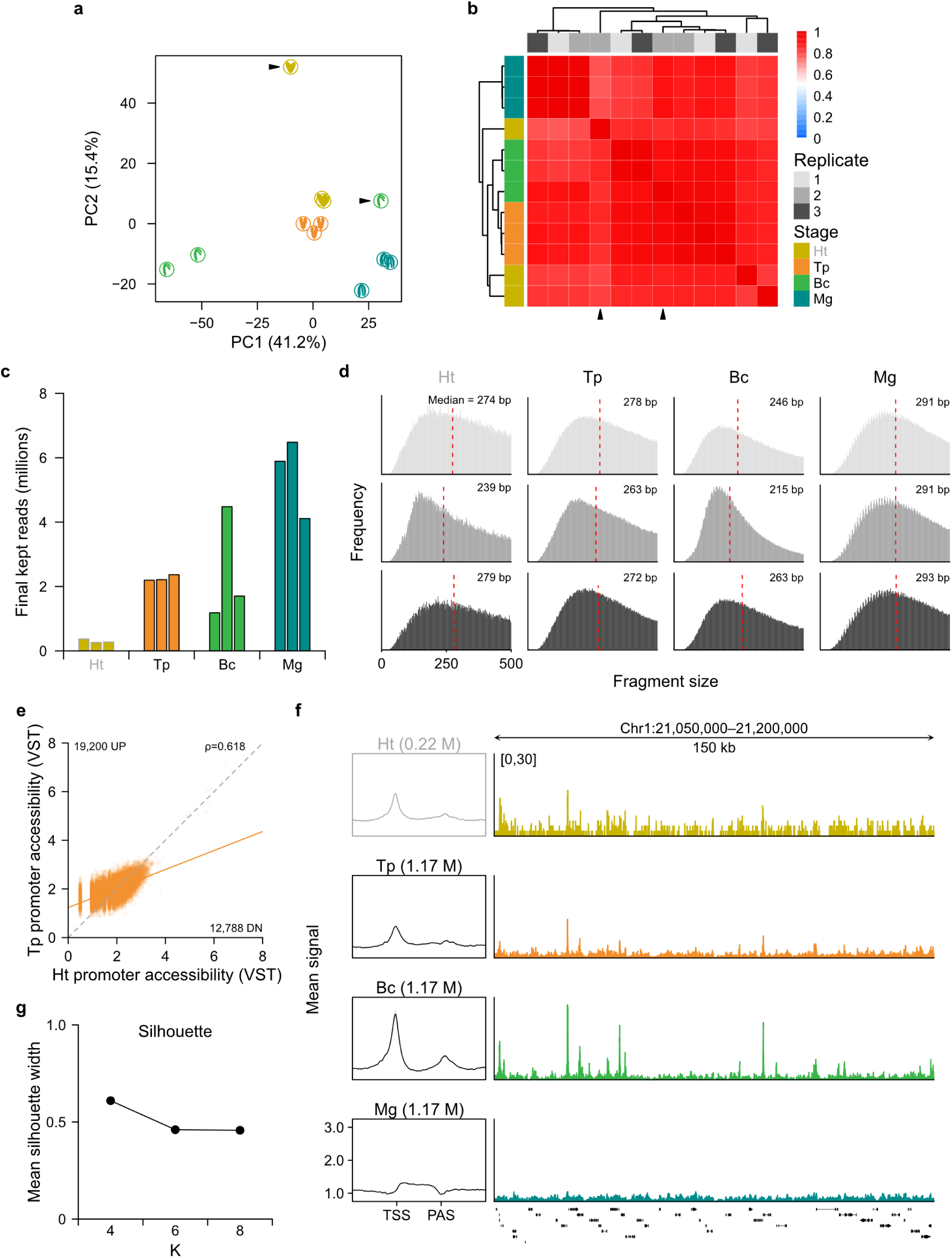
Chromatin-accessibility quality control, sample exclusion, and CA module robustness. **a** PCA of 50-kb binned CA signal across all libraries, including excluded replicates (indicated by arrows). **b** Pairwise Spearman correlation heatmap of 50-kb binned CA signal across all libraries, including excluded replicates (indicated by arrows). **c** Per-library unique read counts across developmental stages and genotypes. **d** Fragment size distributions for each library. **e** Scatter plot of per-gene promoter accessibility for the exploratory T2 transition. **f** TSS- and PAS-centered accessibility metaprofiles after downsampling all core-stage libraries to 1.17 million unique reads, together with representative IGV browser views at selected loci using pooled RPGC-normalized signal. **g** Cluster-quality diagnostics for accessibility module clustering with k = 4, 6, and 8.

**Fig. S3.**
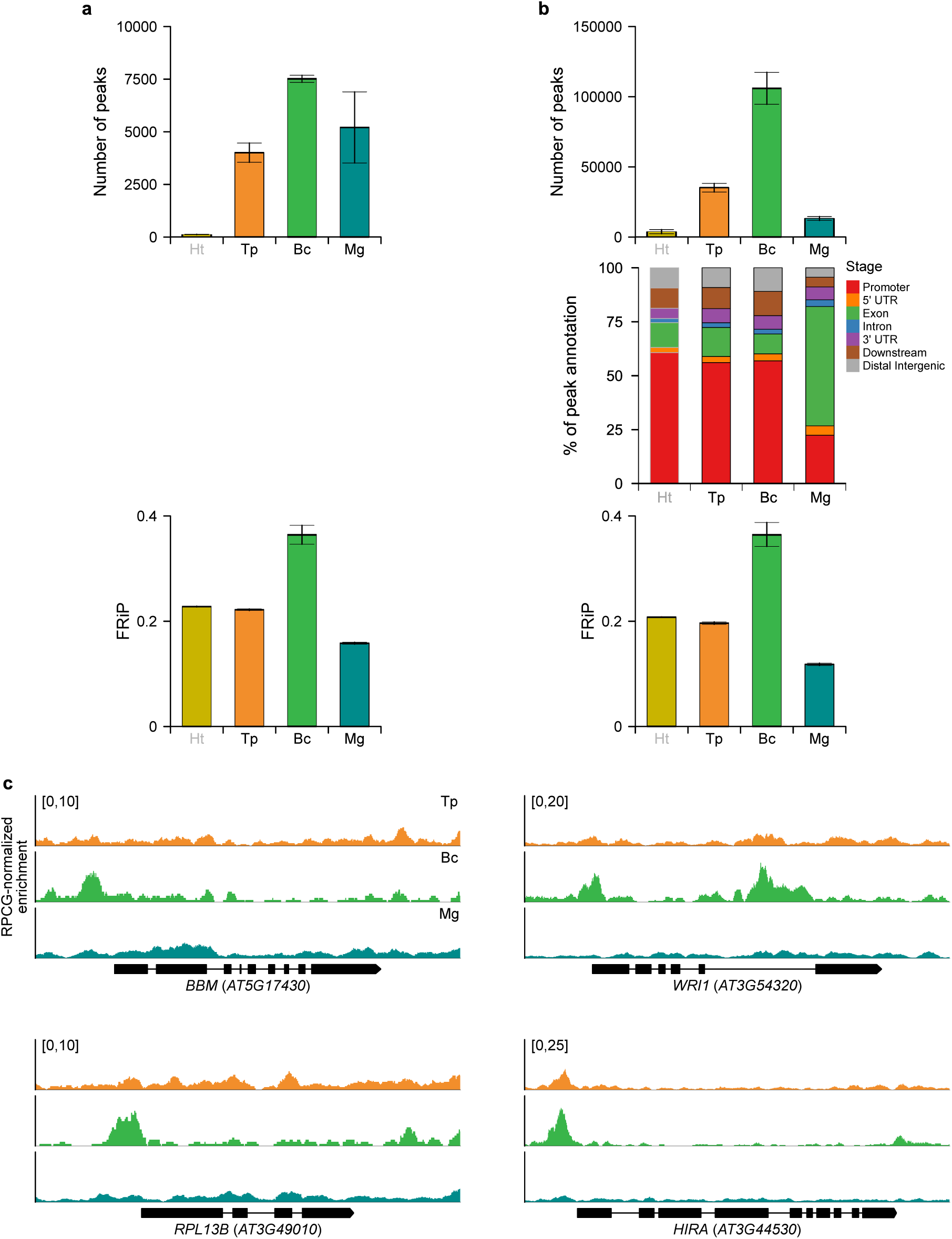
Peak-based accessibility analysis. **a** Number of accessible regions identified by MACS peak calling and corresponding FRiP values across developmental stages. **b** Number of accessible regions identified by window-based peak calling, together with genomic feature annotation and FRiP values. **c** NicE-seq accessibility tracks (pooled RPGC-normalized signal) at four representative loci: BBM, WRI1, RPL13B, and HIRA. Tracks span Tp, Bc, and Mg. Gene models below indicate exon–intron structure and strand orientation. Genomic coordinates extend 1 kb upstream and downstream of each gene body.

**Fig. S4.**
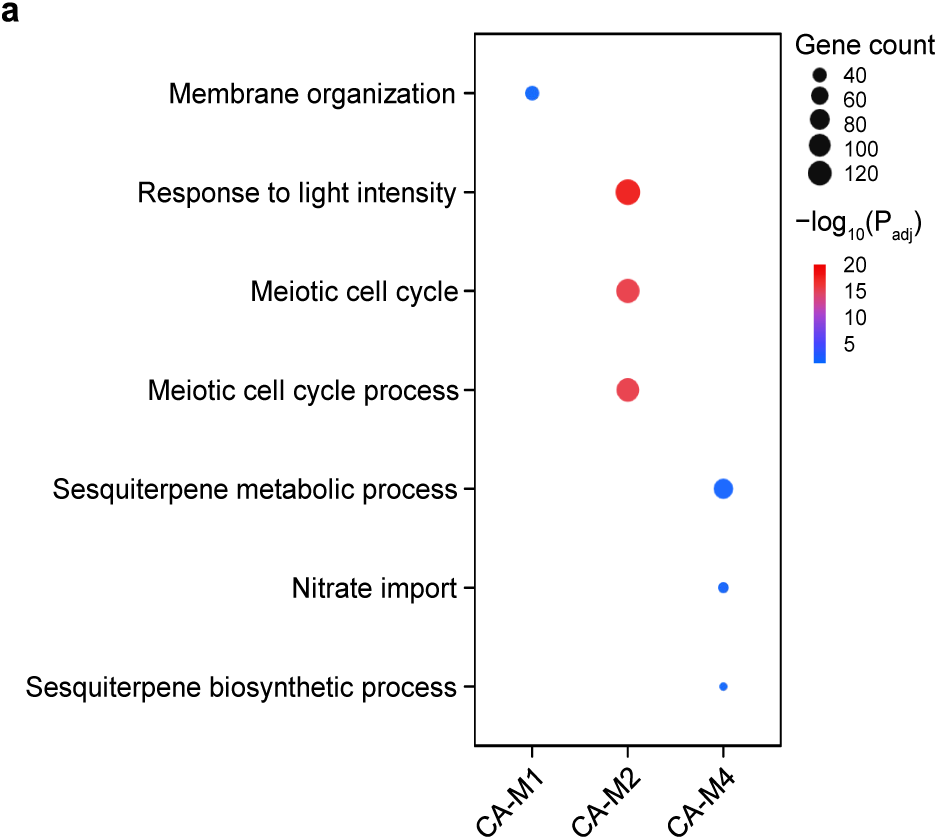
GO enrichment of accessibility modules. **a** GO Biological Process enrichment of accessibility modules (clusterProfiler enrichGO, adjusted P < 0.05). CA-M3 showed no significantly enriched terms and was therefore omitted.

**Fig. S5.**
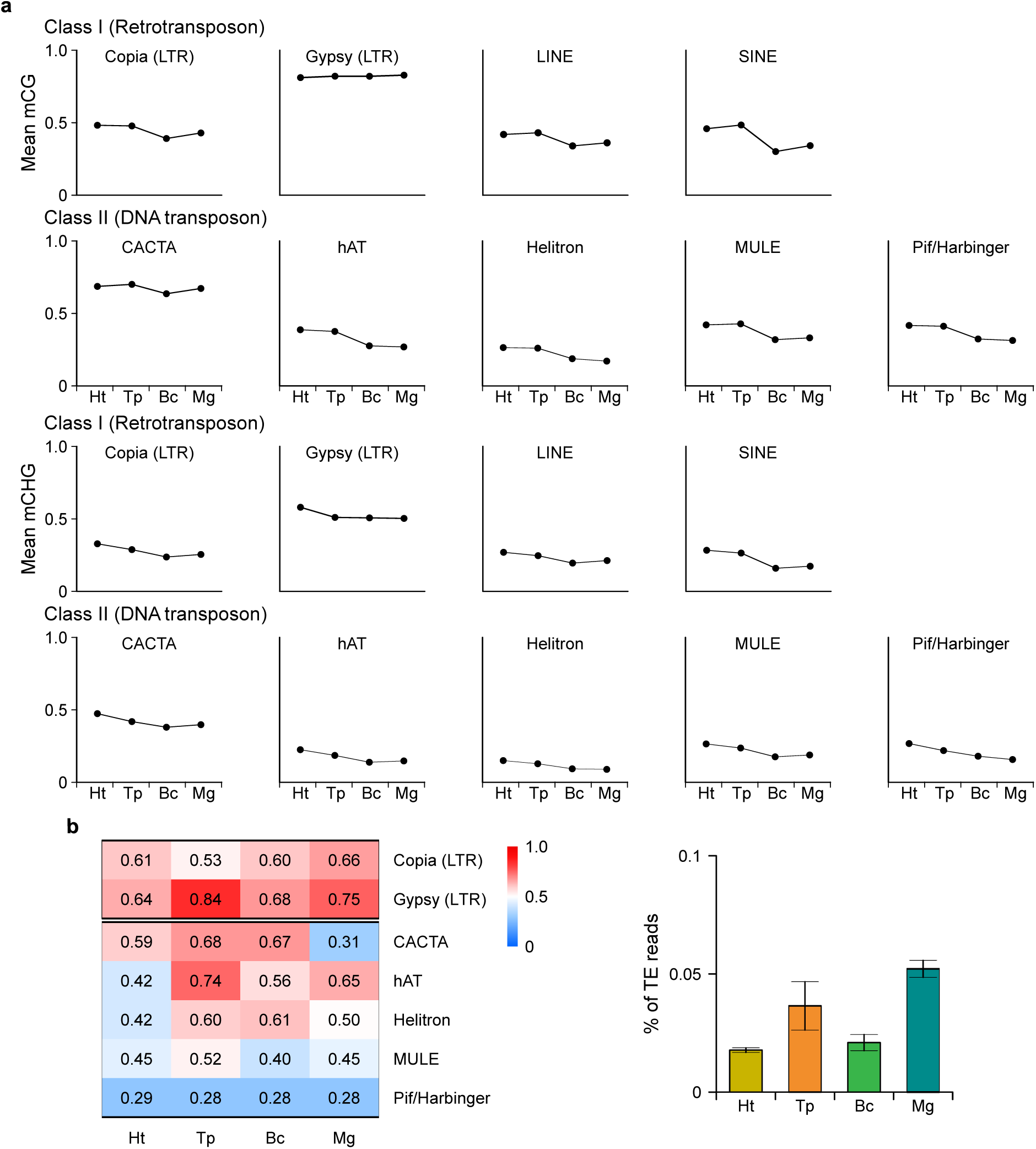
TE body methylation and expression concentration across embryonic stages. **a** CG and CHG methylation levels across TE bodies by TE family and developmental stage, based on re-analysis of public WGBS data (Lee *et al*., 2023). **b** Gini coefficients of TE expression concentration by TE family and developmental stage (left), and fractions of TE-assigned reads among total reads for expressed TE families (right). Gini coefficients were calculated from Nanopore RNA-seq CPM values after exclusion of gene-overlapping reads, whereas TE-assigned read fractions were calculated from read counts after the same filtering step.

**Fig. S6.**
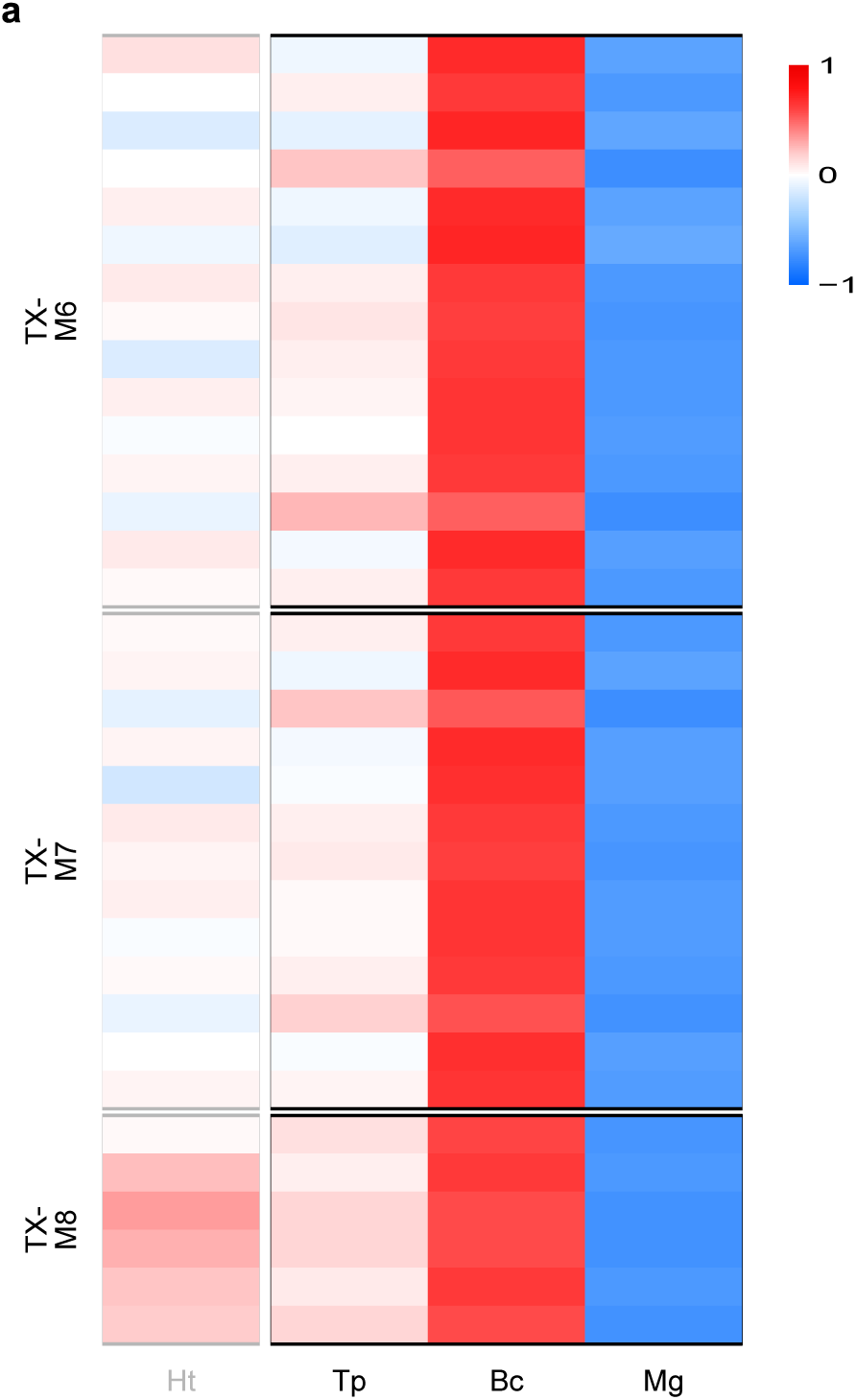
Motif accessibility in late expression modules. **a** Module × TF-family promoter accessibility z-scores for TX-M6, TX-M7, and TX-M8 across Tp, Bc, and Mg, using the same analysis framework as in Fig. 4g.

**Fig. S7.**
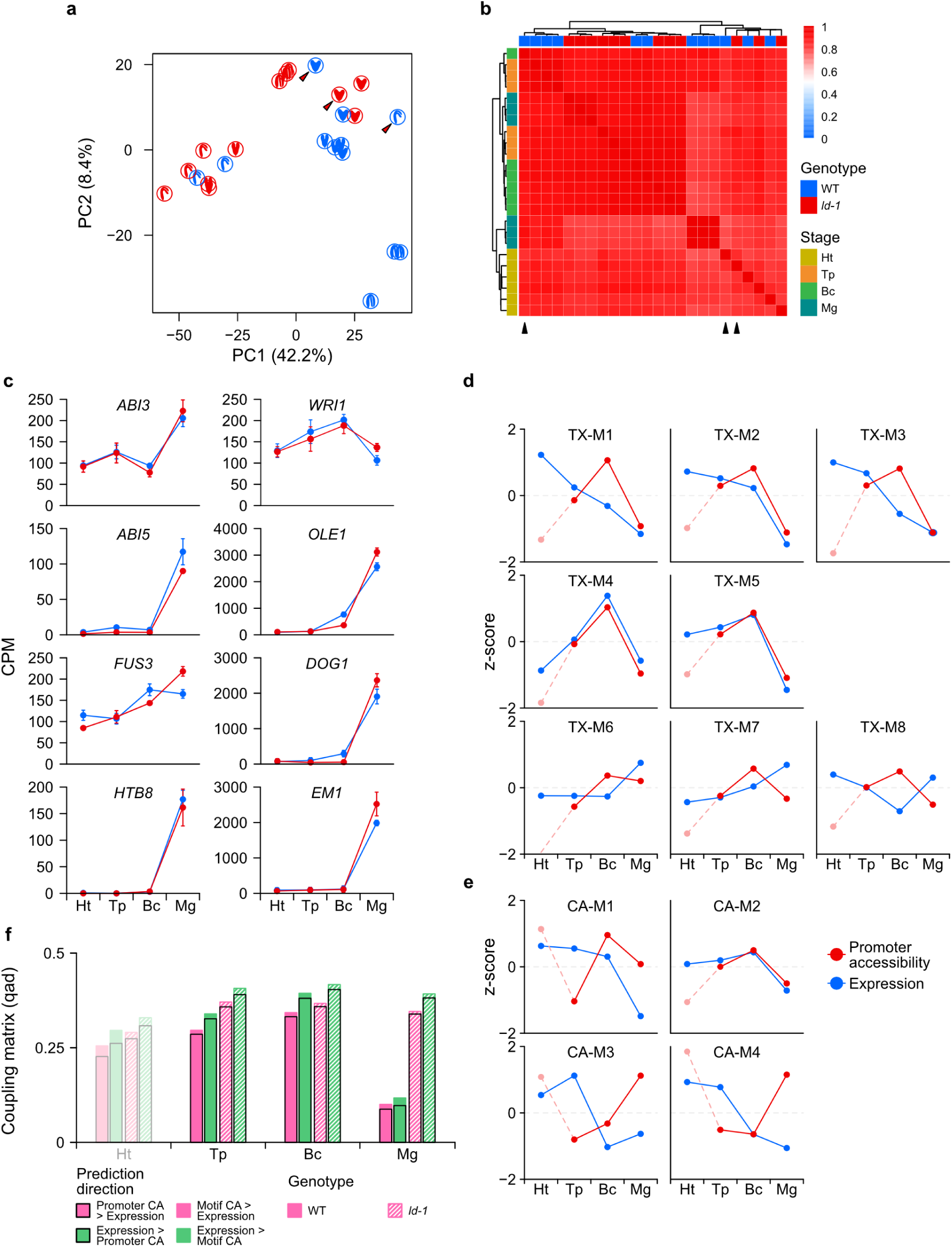
*ld-1* chromatin accessibility sample structure, maturation marker expression, module trajectories, and CA–transcription coupling. **a** PCA of WT and *ld-1* NicE-seq libraries based on 50-kb binned chromatin accessibility signal. Color indicates genotype, and letters indicate developmental stage. Excluded replicates are indicated by arrows. **b** Pairwise Spearman correlation heatmap of WT and *ld-1* NicE-seq libraries based on 50-kb binned chromatin accessibility signal. Genotype and developmental stage annotations are shown above and to the left of the heatmap. Excluded replicates are indicated by arrowheads. **c** Expression trajectories of eight canonical maturation-associated genes: regulators *ABI3*, *ABI5*, and *FUS3*; histone variant *HTB8*; and downstream targets *WRI1*, *OLE1*, *DOG1*, and *EM1*. Values are shown as mean CPM ± SE across developmental stages for WT and *ld-1*. **d** Overlay of expression and promoter accessibility trajectories across WT-defined transcriptome modules (TX-M1 to TX-M8). Expression is shown in blue and promoter accessibility in red. Solid lines indicate WT, and faded dashed lines indicate *ld-1*. Values are displayed as z-scores across developmental stages. Ht is included as an exploratory reference. **e** Overlay of expression and promoter accessibility trajectories across WT-defined accessibility modules (CA-M1 to CA-M4). Expression is shown in blue and promoter accessibility in red. Solid lines indicate WT, and faded dashed lines indicate *ld-1*. Values are displayed as z-scores across developmental stages. Ht is included as an exploratory reference. **f** CA–transcription coupling across developmental stages in WT and *ld-1*, quantified using qad. Coupling was calculated in both prediction directions, from promoter accessibility to expression and from expression to promoter accessibility, and compared with motif-level accessibility–expression coupling shown in Fig. 5h. Ht is ssincluded as an exploratory reference.ss

## Supplementary Tables

**Table S1. Expression module gene lists**

Gene IDs for all eight expression modules (TX-M1 to TX-M8), with each column representing one module.

**Table S2. GO enrichment of expression modules**

GO Biological Process terms enriched in each expression module (clusterProfiler enrichGO, adjusted P < 0.05). Each sheet corresponds to one module and includes gene counts, enrichment ratios, adjusted P values, and associated gene IDs for each term.

**Table S3. Differential expression results**

Pairwise DE results for each consecutive WT stage transition (T2, T3, T4), including log2 fold-change, Wald statistic, and adjusted P values.

**Table S4. GSEA of stage-transition DE genes**

GSEA results for each consecutive WT stage transition (T2, T3, T4). Genes were ranked by the DESeq2 Wald statistic and tested against GO Biological Process terms. Results include normalized enrichment scores (NES), adjusted P value, and leading-edge gene lists.

**Table S5. Accessibility module gene lists**

Gene IDs for all four accessibility modules (CA-M1 to CA-M4), with each column representing one module.

**Table S6. GO enrichment of accessibility modules**

GO Biological Process terms enriched in each accessibility module (clusterProfiler enrichGO, adjusted P < 0.05). Each sheet corresponds to one module and includes gene counts, enrichment ratios, adjusted P values, and associated gene IDs for each term.

**Table S7. Motif enrichment of expression modules**

TF-family-level motif enrichment results for each expression module (monaLisa, JASPAR 2024), including enrichment score, adjusted P value, and the best-matching motif for each TF family.

**Table S8. Genotype-level differential expression results**

Differential expression results for *ld-1* versus WT at each developmental stage, including log2 fold-changes, Wald statistics, and adjusted P values.

**Table S9. Genotype-level GSEA results**

GSEA results for *ld-1* versus WT at each developmental stage. Genes were ranked by the DESeq2 Wald statistic and tested against GO Biological Process terms. Results include normalized enrichment scores (NES), adjusted P values, and leading-edge gene lists.

**Table S10. ChromVAR motif deviation scores**

Per-sample bias-corrected chromVAR deviation z-scores for 810 JASPAR 2024 plant motifs across all retained WT and *ld-1* NicE-seq libraries.

